# Epigenetic Patterns in a Complete Human Genome

**DOI:** 10.1101/2021.05.26.443420

**Authors:** Ariel Gershman, Michael E.G. Sauria, Paul W. Hook, Savannah J. Hoyt, Roham Razaghi, Sergey Koren, Nicolas Altemose, Gina V. Caldas, Mitchell R. Vollger, Glennis A. Logsdon, Arang Rhie, Evan E. Eichler, Michael C. Schatz, Rachel J. O’Neill, Adam M. Phillippy, Karen H. Miga, Winston Timp

**Affiliations:** Department of Biomedical Engineering and Molecular Biology and Genetics, Johns Hopkins University, Baltimore, Maryland, USA; Department of Biology and Computer Science, Johns Hopkins University, Baltimore, Maryland, USA; Institute for Systems Genomics, University of Connecticut, Storrs, CT, USA; Department of Molecular and Cell Biology, University of Connecticut, Storrs, CT, USA; Genome Informatics Section, Computational and Statistical Genomics Branch, National Human Genome Research Institute, National Institutes of Health, Bethesda, MD, USA; Department of Bioengineering, University of California Berkeley, Berkeley, CA, USA; Department of Molecular and Cell Biology, University of California Berkeley, Berkeley CA, USA; Department of Genome Sciences, University of Washington School of Medicine, Seattle, WA, USA; Howard Hughes Medical Institute, University of Washington, Seattle, WA, USA; UC Santa Cruz Genomics Institute, University of California Santa Cruz, Santa Cruz, CA, USA

## Abstract

The completion of the first telomere-to-telomere human genome, T2T-CHM13, enables exploration of the full epigenome, removing limitations previously imposed by the missing reference sequence. Existing epigenetic studies omit unassembled and unmappable genomic regions *(e.g*. centromeres, pericentromeres, acrocentric chromosome arms, subtelomeres, segmental duplications, tandem repeats). Leveraging the new assembly, we were able to measure enrichment of epigenetic marks with short reads using k-mer assisted mapping methods. This granted array-level enrichment information to characterize the epigenetic regulation of these satellite repeats. Using nanopore sequencing data, we generated base level maps of the most complete human methylome ever produced. We examined methylation patterns in satellite DNA and revealed organized patterns of methylation along individual molecules. When exploring the centromeric epigenome, we discovered a distinctive dip in centromere methylation consistent with active sites of kinetochore assembly. Through long-read chromatin accessibility measurements (nanoNOMe) paired to CUT&RUN data, we found the hypomethylated region was extremely inaccessible and paired to CENP-A/B binding. With long-reads we interrogated allele-specific, longrange epigenetic patterns in complex macro-satellite arrays such as those involved in X chromosome inactivation. Using the single molecule measurements we can clustered reads based on methylation status alone distinguishing epigenetically heterogeneous and homogeneous areas. The analysis provides a framework to investigate the most elusive regions of the human genome, applying both long and short-read technology to grant new insights into epigenetic regulation.

## INTRODUCTION

The completion of the original human genome reference in 2001(*1, 2*), has enabled efforts to understand not only coding sequence, but how the rest of the genome acts to regulate gene expression through the epigenome. But the epigenome can only be as comprehensive as the reference it is built on, improving the reference by generating a complete assembly of a human genome (T2T-CHM13)(*3*) provides a unique opportunity to explore the final frontier -- the repetitive areas of the genome.

The vast majority of new sequences in T2T-CHM13 are in the pericentromeric, centromeric, and acrocentric regions (+180.5 Mb) and segmental duplications (+44.2 Mb) which include ampliconic gene families (macro-satellites) in the chromosome arms(*3*). The pericentromeric satellites are made up of human satellites 1-6 (HSat1-6), the beta-satellites (BSat), and the gamma-satellites (GSat). Along with the sub-telomeric satellites, these regions comprise the constitutive heterochromatin domain, playing a critical role in protecting genome integrity(*4*). Epigenetic regulation of the genome controls gene expression but also confers genome stability through regulation of heterochromatin(*5*–*7*). The H3K9me3 histone variant, a mark associated with repressive heterochromatin, is especially prominent in constitutive heterochromatin, contributing to the dense organization and low transcription levels of this region(*8*).

The centromere itself is composed of higher order repeats (HORs), tandem arrays of larger repeat units consisting of multiple basic repeat units. These have a unique epigenetic state, with CENP-A replacing H3 in the nucleosome in centromeric chromatin; epigenetic regulation is thought to play a key role in centromere identity and kinetochore formation(*9*). Localization of CENP-A is not dependent on the underlying DNA sequence, illustrated by the fact that centromeres have been shown to move to new locations along the chromosome, thus encountering new DNA sequences (neo-centomere formation) (*10*). Investigation of histone modifications and variants in repetitive DNA has been precluded by the combination of incomplete human reference assemblies and the reliance on the reduced mappability of short-read sequence data.

Histone modifications often act in tandem with covalent DNA base modifications such as 5-methylcytosine (5mC) at CpG dinucleotides, especially in heterochromatin (*11*). Generation of a high resolution CpG methylation map of satellite regions has been prevented by the combination of incomplete human reference assemblies and the reliance on the reduced mappability of whole-genome bisulfite sequencing (WGBS). Inside these repetitive areas it is especially challenging to examine epigenetic heterogeneity, both along the chromosome and between molecules. Allele-specific methylation(*12*) and epigenetic variation(*13*) can play an important role in gene regulation and transcriptional plasticity, but is hard to study in repetitive areas including regions like DXZ4, a macrosatellite repeat and epigenetic regulator of X chromosome inactivation. In order to probe these questions, precise alignments, anchored to heterozygous genomic elements for allele-specific identification, are typically needed.

Using the T2T-CHM13 complete genome, we have started to explore the epigenome of the newly complete regions. Through the use of k-mer assisted mapping (*14*), we used existing short-read data to explore repeat array-level epigenetic profiles. This high level information provides insight across different cell lines leveraging data from ENCODE(*15*). To look *within* the repetitive areas, we applied long-read epigenetic profiling with nanopore sequencing. Unlike sequencing-by-synthesis strategies, nanopore sequencing directly probes the DNA, allowing for simultaneous measurement of the base sequence and epigenetic state (*16*), with long reads providing deeper insights into the epigenetic patterns on individual molecules. With the combination of the T2T-CHM13 assembly and paired ultra-long-read CpG methylation data from the same CHM13 cell line, we generated the most complete human methylome to date. High resolution methylation maps of centromeric satellites and macro-satellites in the chromosome arms revealed novel mechanistic and epigenetic signatures of these regions, including the epigenetic nature of the human centromere, further validated by simultaneously assessing the complete methylome and chromatin state of the X chromosome centromere in a terminally differentiated, diploid lymphoblastoid cell line (HG002, Coriell GM24385) with ultra-long-read nanoNOMe (*17*). The single-read nature of nanopore sequencing allowed for further insights into the epigenetic cell-to-cell heterogeneity and methylation-based haplotype phasing within nearly homozygous, variant devoid genomic regions. With the combination of massive improvements to complete genome assemblies and the mappability of ultra-long nanopore reads, we show here that investigation of epigenetic regulation in the largely overlooked satellite arrays is both technologically possible and reveals novel mechanistic and regulatory events.

## RESULTS

### Expanding ENCODE Epigenetic Data with Dynamic k-mer Assisted Mapping

The T2T-CHM13 assembly resolves gaps and corrects misassembled or patched regions in GRCh38, leading to nearly 200 Mb of previously unprobed sequence content(*3*). We first examined if we could use existing short-read epigenetic data to gain new insights into the improved T2T-CHM13 assembly. Repetitive DNA regions of particular interest are 1) pericentromeric repeats including the human satellites (HSat1-4), beta-satellites (BSat) and gamma-satellites (GSat) 2) the alpha-satellites which can exist as the kinetochore associated higher order repeat (HOR), a divergent higher order repeat (DHOR) or monomeric sequences (MON) 3) the telomere associated repeats (TAR). These repetitive genomic regions pose issues for short-read alignments. To accurately map short-read epigenetic data to repeat-rich sequences in T2T-CHM13, we developed a dynamic k-mer assisted mapping strategy, where k is dependent on the length of the aligned read (**Fig 1a, Fig S1**, Methods). Our dynamic k-mer assisted mapping strategy differs from previously devised mapping strategies (*14*), in that we account for insertions and deletions (indels) in the reads and change the k-mer size according to the total length of the reference sequence spanned by the read. This is especially important when aligning to cell lines derived from different individuals with expected genetic diversity. As a result, we retain more than 70% of total reads **(Fig S2b)** and can extend k-mer filtered mapping strategies beyond samples where the reads and reference are generated from the same genome (e.g. CHM13).

**Figure 1.**
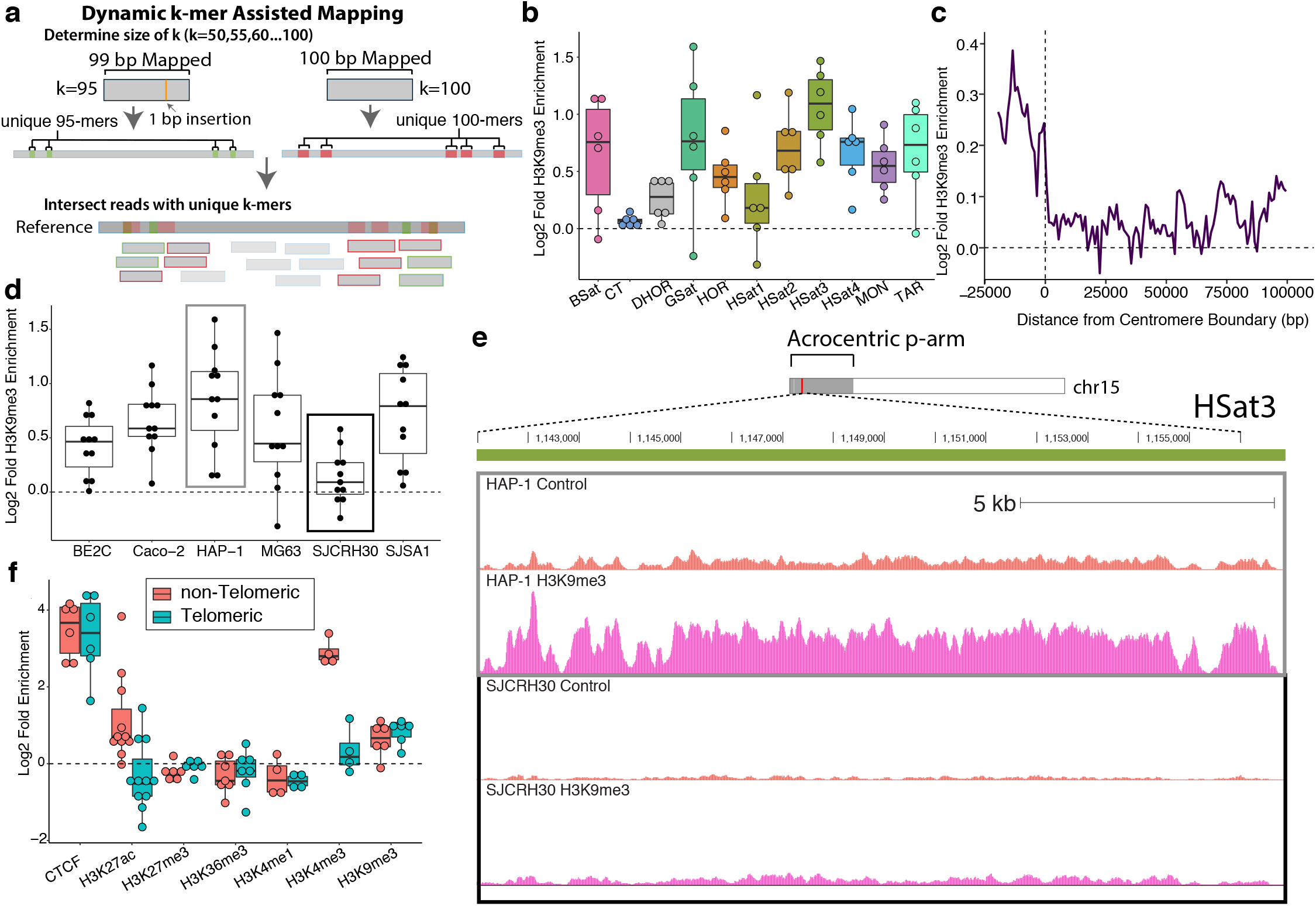
**a)** Schematic of dynamic k-mer assisted mapping pipeline. **b)** Log2 enrichment of H3K9me3 ChIP-seq in satellite repeats normalized to input control. Each point represents an ENCODE cell line. **c)** Log2 fold enrichment of H3K9me3 at the centromeric edges. Negative x-coordinates are within the satellite array and positive x-coordinates are the number of base pairs past the end of the last centromeric satellite array. **d)** Log2 enrichment of H3K9me3 ChIP-seq normalized to input control for each satellite repeat. Each dot represents a satellite repeat class: HSat1, HSat2, HSat3, HSat4, GSat, HOR, DHOR, BSat, MON, TAR and CT. **e)** UCSC genome browser tracks of H3K9me3 coverage versus input control (no antibody) for HAP-1 (highest enrichment) and SJCRH30 (lowest enrichment). **f)** Log2 fold enrichment of different epigenetic marks in the telomeric TAR repeats (within 2 kb of chromosome end) versus the non-telomeric TAR repeats.

We applied dynamic k-mer assisted mapping to ChIP-seq datasets generated as part of the ENCODE project (*18*). Across six different histone marks and CTCF, an important regulator of chromatin architecture, on average 2.35% more reads mapped to T2T-CHM13 than GRCh38 (**Fig S2, Table S1, Supplemental Data 1)**. Reads filtered by unique k-mer intersection from GRCh38 were enriched by as much as 35% and averaging 6.5% in satellite regions, LINEs and SINEs, segmental duplications, and in general across the genome (**Fig S3**). All centromeric, pericentromeric, and subtelomeric repeats were enriched in H3K9me3, with only the subtelomeric Telomere Associated Repeat (TAR) enriched for activating histone marks **(Fig 1b, Fig S4)**. Examining the H3K9me3 enrichment of the centromeric and pericentromeric satellites, the alpha satellite HOR, HSat2, and HSat3 were more enriched than HSat1 **(Fig 1b)**. Interrogating pericentromeric boundaries, here defined as the position of the last annotated satellite repeat flanked by unique sequence, we observed these boundaries were sharply demarcated by a transition from constitutive heterochromatin as measured by H3K9me3. This is consistent with the heterochromatin domain boundaries observed in yeast and drosophila (*19*–*21*), and indicates the lack of a gradual transition from heterochromatin to euchromatin **(Fig 1c)**.

Next, we used this method to clearly distinguish samples that have heterochromatin dysfunction, which is known to have biological consequences (*4*). While all six cell lines have an enrichment of H3K9me3 in satellite DNA, SJCRH30, a rhabdomyosarcoma derived cell line, has overall lower H3K9me3 enrichment in all satellite repeats, potentially indicating satellite epigenetic dysregulation as a clinically relevant pathology in rhabdomyosarcoma **(Fig 1d)**. This trend can be observed with more detail in an HSat3 repeat on the acrocentric arm of chromosome 15, where H3K9me3 in SJCRH30 is clearly depleted in comparison to HAP-1 **(Fig 1e)**

In contrast to the heterochromatic H3K9me3, we found increased alignments in activating marks; H3K4me3 (2.47%), a distinctive mark of active or poised promoters (*22*), H3K27ac (2.33%), an active enhancer mark (*23*), and CTCF (2.51%), a chromatin binding factor (*24*). Many of these marks fall within the TAR sequence, located 2 kb upstream from the canonical telomeric repeat on all T2T-CHM13 chromosome arms except 8q, 13p, 14p, 16p, 17p, 21p, 22p, and Xp. A CTCF site in the TAR loci drives transcription of the TERRA lncRNA (*25*); a negative regulator of telomerase-mediated telomere elongation. We observed enrichment of CTCF in all ENCODE cell lines at the TAR loci **(Fig S4)**.

Additionally, the subtelomeric regions are rich in segmental duplications resulting in the TAR sequence being dispersed throughout the genome (*26*). When comparing telomeric TAR sequences to non-telomeric TAR sequences we do not observe statistically significant differences (Kruskal-Wallis, p-value=0.12) in sequence divergence (**Fig S5)**. While both telomeric and non-telomeric TAR sequences are enriched for CTCF, the non-telomeric TAR sequences are more enriched for activating chromatin marks H3K27ac and H3K4me3, suggesting differences in TERRA activity **(Fig 1f**). Previous studies have focused on the roles and regulation of TAR in the subtelomere (*27*–*29*), however our results indicate biological functionality of TAR sequences outside the subtelomere as well. Overall, using only publically available short-read data with the new T2T-CHM13 reference and dynamic k*-*mer assisted mapping, we uncovered novel regulatory features in centromeric, pericentromeric, and subtelomeric repeats and display the usage of this technique for investigating disease related epigenetic dysregulation.

### DNA Methylation Measurement in Satellite Repeats with Nanopore Sequencing

Short reads can provide large-scale enrichment profiles of chromatin modifications and DNA binding proteins, but it is especially difficult to probe DNA methylation in repetitive sequences due to mapping biases after bisulfite treatment (**Fig S6)** (*30*). But a significant portion of the CpGs in the genome are in these difficult to probe highly repetitive regions and their epigenetic dysregulation can result in cancer, irregular transcription of regulatory RNAs, and improper chromosome segregation (*31*–*33*). In the T2T-CHM13 assembly, we identified 32.3 million CpG sites, a 9.04% increase from the 29.2 million identified in GRCh38 (omitting chromosome Y) **(Fig 2a, Table S2)**. Overall, in T2T-CHM13, we detected an additional 1.65 million CpG sites in satellite DNA, increasing the percentage of CpGs in satellite DNA from 3.6% in GRCh38 to 8.3% in T2T-CHM13 **(Fig 2a, Table S2)**. Some of the regions have fewer CpGs in CHM13 than in GRCh38 (**Fig S7**). We attribute this to the patchwork nature of GRCh38 being composed of multiple individuals (*34*).

**Figure 2.**
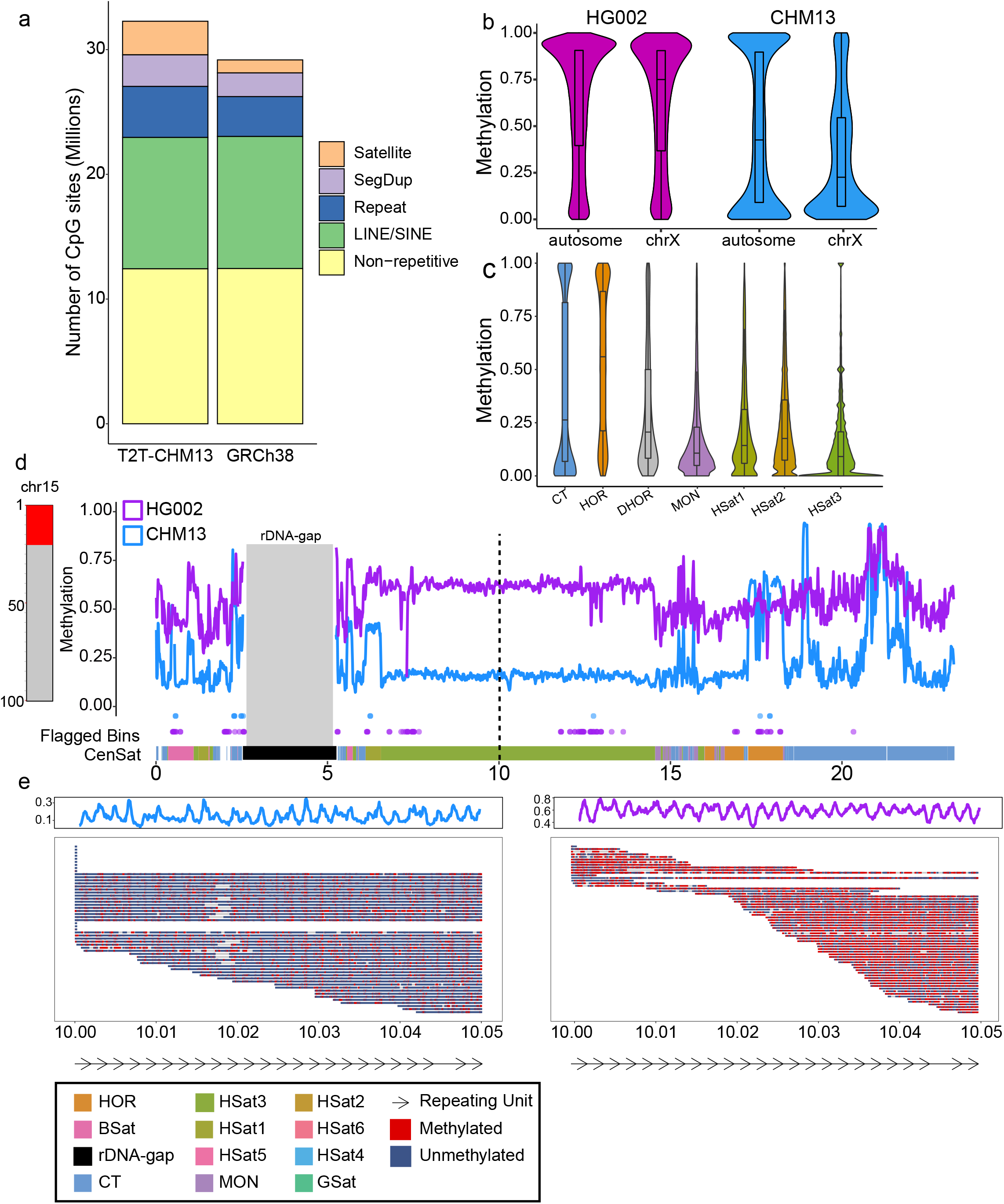
**a)** Percentage of CpG sites contained within different genomic regions (Methods) **b)** Methylation frequency distribution for CHM13 and HG002 non-centromeric regions. **c)** Per-base methylation frequency distributions within CHM13 satellite repeats. CpG site density for corresponding repeat regions shown in **Fig S13. d) (Left)** Vertical chromosome ideogram represents the entirety of chromosome 15 with red representing the acrocentric region plotted. **(Right)** Methylation frequency panel of the acrocentric region of chromo-some 15. Smoothed methylation in 10 kb bins (Methods) is plotted for both CHM13 (blue) and HG002 (purple). The rDNA region in the T2T-CHM13 assembly is annotated by the grey rectangle, methylation calls were not generated in rDNA arrays as these are artificially homogenized model sequences. Blue and purple dots under the methylation frequency represent flagged bins with anomalous coverage. Bottom bar shows annotations of satellite repeats. **e)** Single-read methylation analysis of reads greater than 50 kb in the center of the HSat3 repeat array. Location of region is denoted in **d)** by the vertical dashed line. Each row represents an individual nanopore sequencing read. Red CpG sites are methylated and blue unmethylated. Summary smoothed frequency for the region is shown above the single-read data. Arrows below plots annotate the 1.8 kb repeating units of the HSat3.

We used long-read nanopore data with nanopolish to characterize methylation in the satellite repeats(*16*). Using reads longer than 50 kb greatly improved our ability to map to centromeric and pericentromeric regions, evidenced by declining coverage biases as read length increases (**Fig S8a-b**). But nanopore sequencing has different challenges than WGBS; both general basecalling quality and the quality of methylation calls can be influenced by low-complexity regions which may decrease the signal-to-noise ratio as 6-mers with small shifts in current pass through the pore(*35, 36*). For nanopore methylation analysis, we used quality cutoffs (Methods) to identify a set of high quality methylation calls (71.72% of all methylation calls), where one CpG on one read represents a single call (**Fig S8c**). We observed only a slight decrease in the proportion of high quality calls in the human satellite repeats versus the rest of the genome (**Fig S9a-b, Table S3**). The lowest quality calls were in HSat3 and HSat4, where the percentage of high-quality calls were 62.47% and 65.51%, respectively, compared to the whole genome of 71.72%. Because of the decreased 6*-*mer diversity in HSat repeats, we were concerned that HSats were enriched for 6-mers with small current shifts from methylation, precluding high quality calls. To inspect this, we extracted 6-mers from each HSat repeat array and quantified the average current shift in the nanopore raw data due to methylation. The only notable difference was a *larger* methylation-induced current shift associated with k-mers from HSat2 **(Fig S9c)**, suggesting the repetitive sequence does not generate any specific challenges to methylation calling. Furthermore, we observe low correlation (R=.36) between the mean current shift and the percentage of high quality calls per satellite repeat class **(Fig S9d)**. However, there appeared to be a slightly stronger relationship (R=.42) between satellite class read coverage and percent high quality calls **(Fig S9d)**, suggesting mappability plays a larger role in classification of methylation signal in satellites than delineation of current signal. Finally, we investigated strand bias in methylation calls and noted higher negative strand coverage in HSat4 and HSat5 and higher positive strand coverage in HSat2, but these slight differences in strand specific mappings did not influence methylation frequency results (**Fig S9e-f**). From this analysis, we can be confident that our methylation calls from nanopore sequencing are accurate in satellite regions.

### Long Reads Generate Complete Methylation Maps of Human Chromosomes

The CHM13 cell line is derived from a complete hydatidiform mole, a zygote that develops with two copies of paternal chromosomes and loss of maternal chromosomes. This uniparental disomy should result in exclusively paternal imprinting coupled to global genome demethylation (*37, 38*). Comparing CHM13’s methylation state to existing DNA reduced representation bisulfite sequencing data on early human embryos (**Fig S10, Table S4**) (*39*) we observed that CHM13 clusters closely with cleavage and blastocyst-stage embryos as well as trophectoderm tissue.

To compare the epigenome of the early developmental stage CHM13 cell line to a terminally-differentiated adult cell line, we performed nanopore sequencing of nucleosome occupancy and methylome (nanoNOMe) on HG002, a male lymphoblast line. Of the 32.3 million CpG sites in the CHM13 reference assembly, 99.37% of CpG sites in CHM13 reads and 98.54% of CpG sites in HG002 reads contained a high-quality methylation call. The difference in the percentage of called sites between the two cell lines is a result of the superior mapping of CHM13 to its own genome. The matched sequencing reads and reference assembly on CHM13 reduce genomic variants, differences in array structure, and repeat copy number variation between the reads and assembly. CHM13 also represents only a single haplotype mapping to a haploid representation of the genome, while HG002 is diploid, further complicating alignment to highly variable repetitive DNA (*40, 41*).

To set a baseline avoiding any mappability bias, we compared global methylation distribution between the non-centromeric regions; as expected, CHM13 (36.8% median methylation) was globally hypomethylated compared to HG002 (75% median methylation) **(Fig 2b)**. HG002 has the typical methylation pattern observed in terminally differentiated cells where the majority of the genome is 75% methylated with a secondary peak of unmethylated regions largely represented by CpG islands. In contrast, CHM13 is dramatically hypomethylated as expected from a trophoblastic cell line (*39, 42, 43*). Looking more closely into mapping of nanopore epigenetic data to the satellite regions assembled from T2T-CHM13, we identified regions of coverage dropout and pile-up by flagging CpG loci with coverage lower than 10X and greater than 100X for both HG002 reads and CHM13 reads mapped to the T2T-CHM13 reference (**Fig S11**). These regions only accounted for 1.1% of total sites in CHM13 and 4.26% in HG002, however, a disproportionate amount occurred within the HSat3 array of chromosome 9, (69.10% in CHM13 and 22.93% in HG002). This can be mostly attributed to mapping difficulties; the HSat3 of chromosome 9 is part of the curated list of remaining anomalous regions(*3*). The HOR, HSat1, and HSat2 satellite regions had significant increases in coverage biases in HG002 while other multi-megabase satellite arrays, such as the HSat3 on chromosome 15, had more regular coverage profiles **(Table S5, Fig S12)**.

Examining a specific region, we interrogated the short arm of chromosome 15; previously unassembled in GRCh38 due to its highly repetitive sequence content. The improved T2T-CHM13 assembly and satellite DNA annotation revealed more than 25 Mb of sequence, 7.9 Mb of which is a HSat3 nested tandem satellite repeat(*44*)**(Fig 2d)**. Using long reads we characterized methylation patterns within the highly repetitive HSat3 tandem repeat at even greater detail. We plotted single-read epigenetic data in a 50 kb region at the center of the chromosome 15 HSat3 array **(Fig 2e)**. This allowed us to identify repeating methylation patterns consistent with the repeating 1.8 kb HSat3 sequence (**Fig 2e**; arrows). We note that though CHM13 is hypomethylated as compared to HG002, the periodic fluctuation in methylation signal is consistent in both cell lines. In the CHM13 data, we observe a 1.8 kb deletion 26.250 26.275 26.300 26.325 Mb (chr15:10,017,199-10,019,076) present in a fraction of the reads which corresponds to the deletion of a single HSat3 repeat unit on one haplotype, highlighting the power of long-read sequencing in providing simultaneous information on genetic and epigenetic state.

### Methylation Maps of Alpha-Satellite Repeats Reveal Complex Epigenetic Patterns

Using the superior mapping we obtain from the CHM13 nanopore data to its own genome, we characterized methylation in centromeres and pericentromeres. We found that in this early developmental cell line, all of the satellites showed low methylation levels with the exception of the alpha-satellite HOR arrays **(Fig 2c)**. Notably, we identified distinct regions of hypomethylation within the otherwise hypermethylated centromeric arrays of all CHM13 centromeres, which we term the Centromeric Dip Region (CDR) (**Fig 3**) (*14, 45*). The CDR only occurs within “live” HOR arrays (i.e. the functional unit of the centromere associated with kinetochore attachment) and not within “dead” HOR arrays, which are remnants of the centromeres of human pre-great ape ancestors (*44, 46*)(**Fig 3a, Fig S14**). The HOR hypermethylation and presence of the CDR in the “live” HOR array is consistent across every CHM13 centromere (**Fig S14)**, suggesting that it is of functional significance to the proper maintenance and activity of the centromere. We observed that although chromosome 3 has multiple live arrays broken up by an HSat1 array, it has only one CDR **(Fig 3a)**, suggesting that even between live arrays, the CDR demarks a functionally active site.

**Figure 3.**
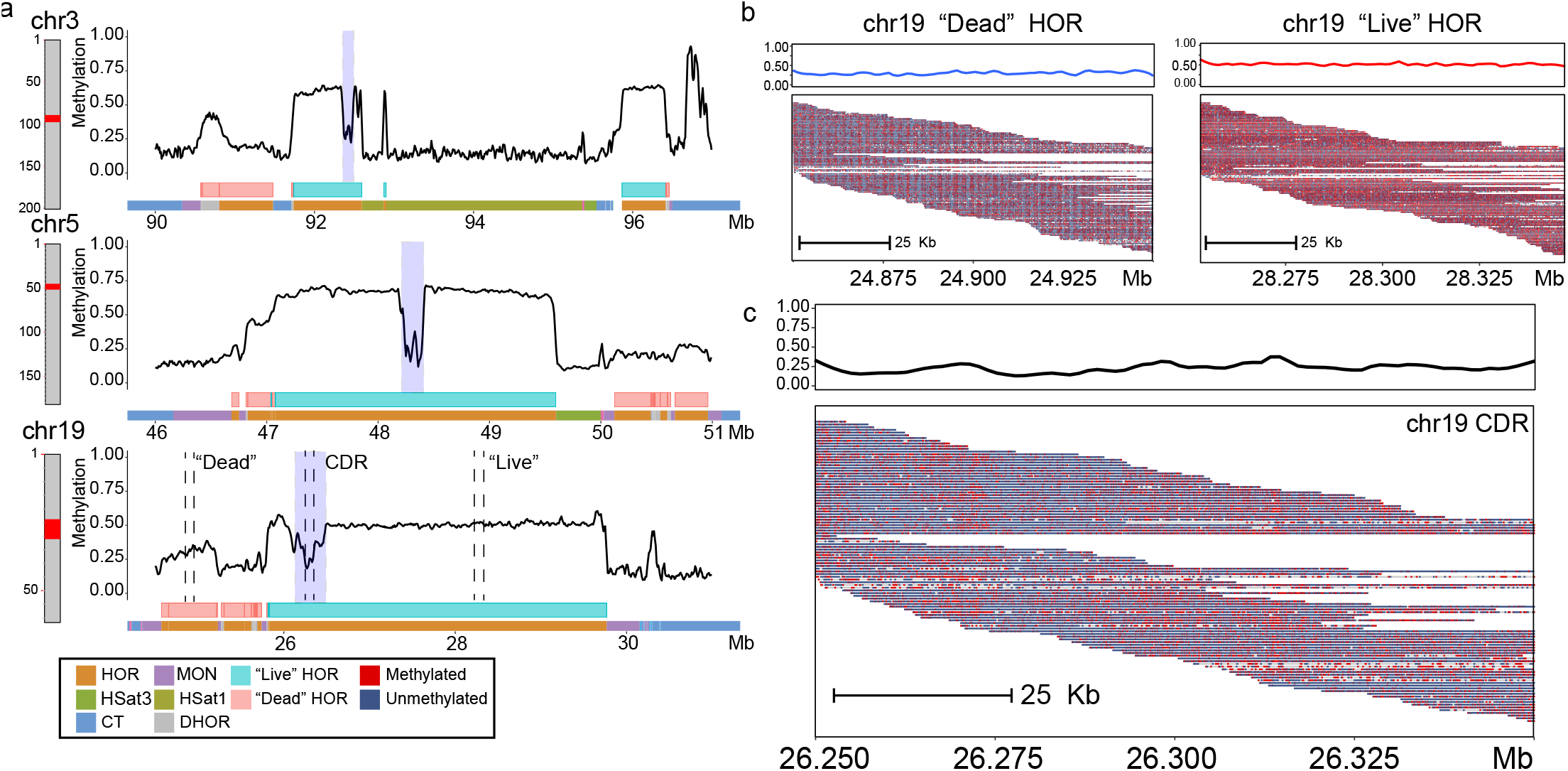
Methylation signatures in HORs are consistent across all CHM13 centromeres. **a)** Panels representing the centromeric regions of chromosomes 3, 5, and 19. Methylation frequency is plotted as smoothed methylation in 10 kb bins (Methods). Pink and grey boxes annotate the “live” and “dead” HOR arrays. Chromosome ideogram is colored with satellite repeat annotations. **b)** Single-read alignments comparing the “live” and “dead” HOR arrays on chromosome 19. Red line is the average methylation frequency of the “live” HOR and the blue line is average methylation frequency of the “dead” HOR fitted with a LOESS model. **c)** Single-read alignments within the CDR on chromosome 19. Black line is average methylation frequency fitted with a LOESS model.

Though not all chromosomes contain “dead” HOR arrays, chromosomes 3, 5 and 19 do have prominent dead arrays **(Fig 3a, Fig S14)**. The dead HOR arrays have a median methylation of 22.6% as compared to the live array median of 60%. From the single-read plots we observe a methylation banding pattern on the dead HOR array that is not present in the live array. The banded hypomethylation suggests a higher-order epigenetic structure within the dead HOR array within the repeating alpha satellite units **(Fig 3b)**. Within the CDR region, we observed a pattern defined by large regions of hypomethylation and periodic increases in methylation **(Fig 3c)**.

### Conservation of Alpha-Satellite Epigenetics in Terminally Differentiated Diploid Cell Lines

To investigate if the CDR also appears in terminally differentiated adult cells, we examined the HOR in HG002. To alleviate the difficulties involved in alignment to this highly divergent region, we aligned to a complete assembly of the HG002 chromosome X centromere(*3*). With our nanoNOMe data accurately aligned to this region, we measured both chromatin accessibility and methylation, using reads greater than 20 kb to achieve a sequencing depth of 40X (**Fig S15f**). In this method, we use M.CviPI methyltransferase to decorate accessible chromatin with exogenous GpC methylation (*17*); using GpC sites we can then measure chromatin accessibility along with CpG methylation of the HG002 sequencing reads aligned to the HG002 chromosome X centromere assembly. Examining methylation we clearly observe a CDR in the HG002 chromosome X HOR region (**Fig 4a,b)**. The CHM13 HOR median methylation was similar to the HOR median methylation in HG002 chrX (56% in CHM13 and 65% in HG002 chrX) (**Table S6-7**). Outside the CDR, the HOR is more hypermethylated in HG002 (65% median methylation) than in CHM13 (30.8% median methylation), consistent with the overall hypomethylation of the X chromosome in CHM13.

**Figure 4.**
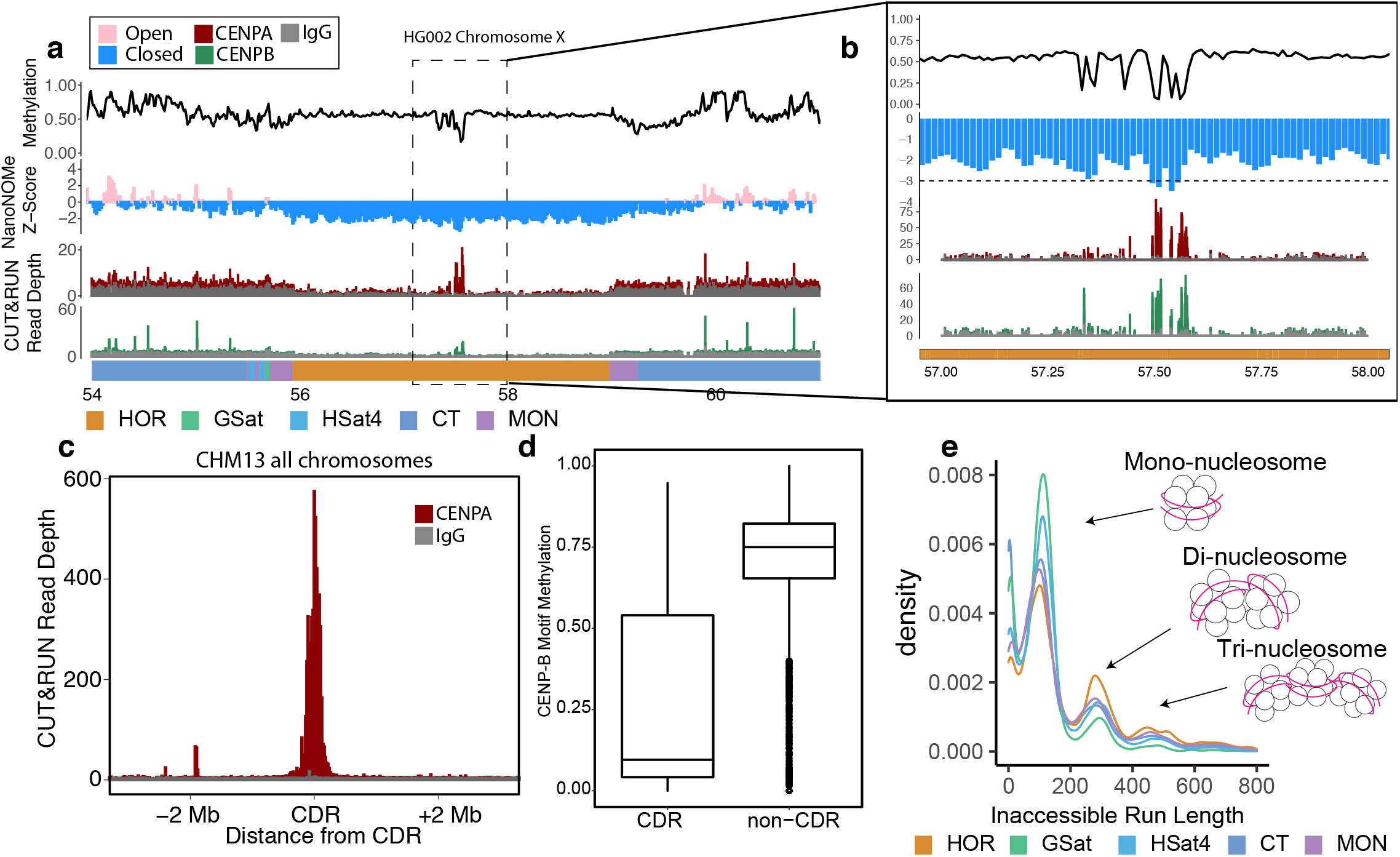
HOR hypomethylation is associated with inaccessibility and CENP-A. **a)** Methylation, nanoNOMe accessibility, CENP-A and CENP-B CUT&RUN data across the chromosome X centromeric array on HG002. Smoothed methylation and accessibility are plotted in 15 kb bins (Methods), CUT&RUN is plotted as raw read counts. Bottom bar shows satellite annotations indicating the location of the HOR, GSat, HSat4 and centromeric transition flanking regions. **b)** Zoom of **a)** in the CDR region. Dotted line provided as guide to eye at Z-score of -3. **c)** Average CENP-A CUT&RUN coverage relative to the CDR position on all CHM13 chromosomes. **d)** Methylation frequency of the CpG sites within the CENP-B motif inside and outside the CDR (p<1e-15) on HG002 chromosome X. **e)** Density plots of length of inaccessible run lengths within different genomic regions on HG002 chromosome X.

To measure the chromatin state relative to the entire chromosome, we calculated GpC methylation Z-scores throughout HG002 chromosome X in 15 kb bins for which the bin size was chosen to avoid bias in distribution of GpC motifs (**Fig S15a-g)**. Bins with lower Z-scores (negative) are more inaccessible and bins with higher Z-scores (positive) more accessible **(Fig 4a**,**b, Fig S15e)**. Remarkably, the most inaccessible region of the entire X chromosome was at 57.49-57.57 Mb, corresponding to the CDR. We hypothesized that this region is inaccessible due to blockage of the GpC methyltransferase by the kinetochore complex and higher-order structure of CENP-A chromatin (*47*), a centromere associated histone variant that directs the assembly of active kinetochores (*48*).

To test our hypothesis that the inaccessibility at the CDR is due to CENP-A chromatin, we analyzed CHM13 CENP-A CUT&RUN data as well as HG002 CENP-A and CENP-B CUT&RUN data using marker-assisted-mapping(*44*). From a metaplot of CDRs across all chromosomes in CHM13, we see a strong enrichment of CENP-A at the CDR compared to the IgG control (**Fig 4c**). In HG002, we see similar enrichment of CENP-A and CENP-B with prominent peaks in the CDR region **(Fig 4a)**. We also noted a marked decrease in coverage from the IgG control across the HOR array as compared to the centromeric transition regions, implying a decline in chromatin accessibility **(Fig 4a)**. Assessing the coordination between methylation and CENP-B motifs, we found a significant difference (p<1e-15) in methylation between motifs within the CDR (9.5% median methylation) and outside the CDR (75% median methylation); consistent with *in vitro* studies showing the preference of CENP-B in binding an unmethylated CENP-B box (**Fig 4d)** (*49*). Zooming in on the CDR, we identified hypomethylated valleys that overlap with the nanoNOMe peaks and CENP-A peaks (**Fig 4b**), potentially indicating kinetochore attachment sites and further validating the relationship between the hypomethylation and centromere protein localization.

Another advantage of profiling accessibility with long nanopore reads is that we can compare nucleosome density within satellite arrays in the centromeric regions of HG002 chromosome X **(Fig 4e)**. By assessing the length of inaccessible (GpC unmethylated) runs, we can find lengths corresponding to mono-nucleseosome (130 bp), di-nucleosome (260 bp) and tri-nucleosome (420 bp) peaks. Though clearly visible when looking across all of chromosome X, we do not see these inaccessible runs in reads from the histone-free mitochondrial genome, indicating efficient chromatin labelling **(Fig S16)**. We compared the mono-, di-, and tri-nucleosome content between different centromeric satellite repeats in HG002 chromosome X, namely the HSat4, MON, GSat, HOR and for comparison, the CT. **(Fig 4e)** (*17*). The HOR array has the highest proportion of di- and tri-nucleosomes comparatively, indicating it is the most nucleosome dense region of the centromere (**Fig 4e**). The GSat array has a high proportion of mono-nucleosomes when compared to the other centromeric and pericentromeric regions; this more euchromatic state is consistent with previous literature (*50*).

### Single Molecule Epigenetics Reveals Haplotype Specific Regulation

The long reads, coupled to a complete reference assembly, confer the ability to explore methylation patterns of single molecules, each of which represents the methylation pattern of a single allele from a single cell. Using this we were interested in visualizing epigenetic heterogeneity within the CDR regions. We focused on the only haploid centromere we profiled, the HG002 chromosome X CDR, to isolate allele-specific effects from cell-to-cell epigenetic heterogeneity. When organizing reads at the distal edge of the CDR by their average methylation, we noticed significant heterogeneity at the edges of the CDR in HG002, showing cell-to-cell variability in the region boundaries **(Fig 5a)**. This variability suggests dynamic regulation of methylation in the CDR similar to what is observed in 5mC border encroachment of CpG islands in cancer (*51*). However, as the reads approach the center of the CDR, the region is consistently unmethylated **(Fig 5a)**.

**Figure 5.**
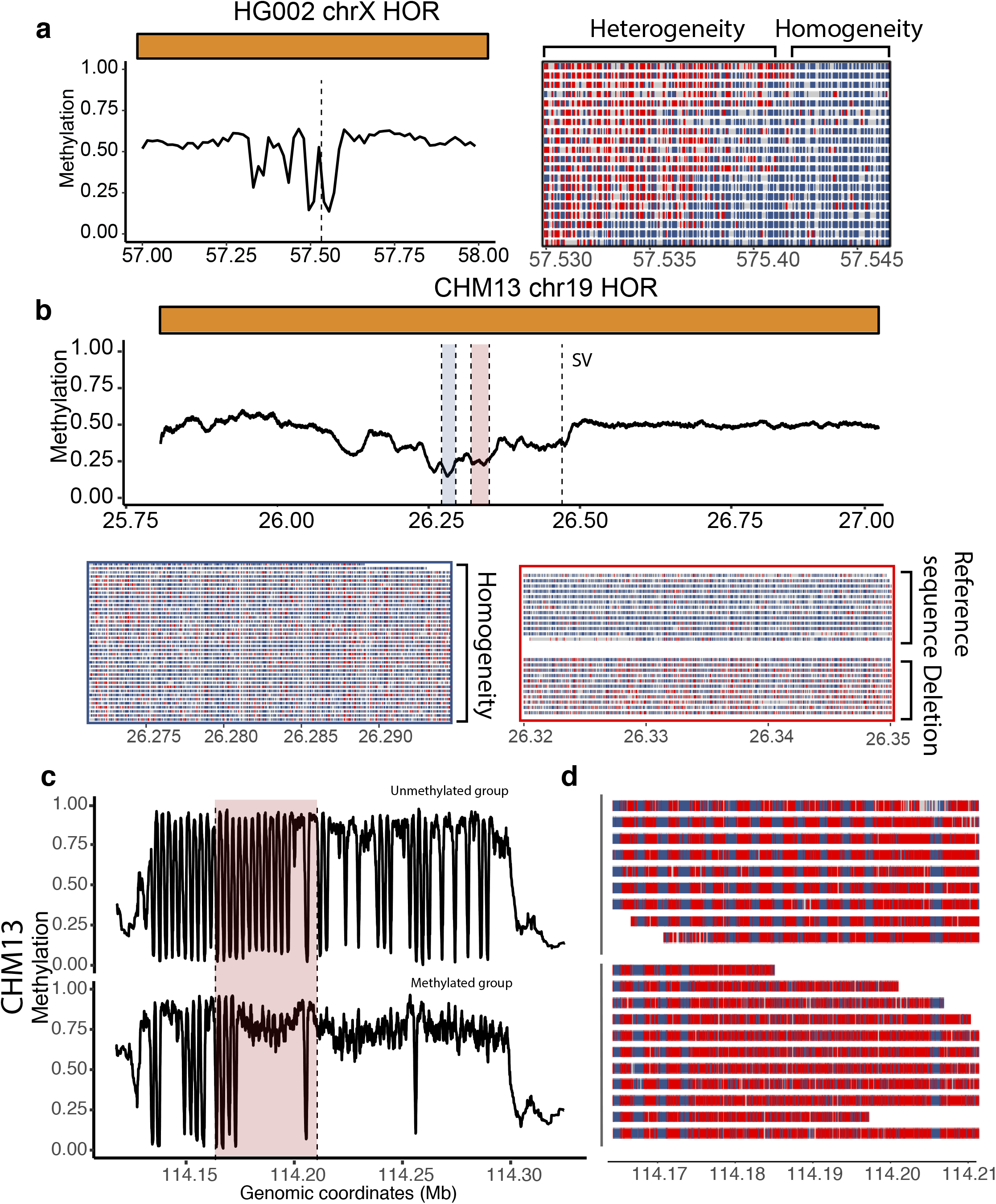
Single molecule Methylation Patterns. **a)** (**Left)** Methylation frequency in the CDR region of HG002 chromosome X. **(Right)** single-read methylation calls in the region indicated by the dashed black line. **b) (Top)** the CDR region of chromosome 19 with the homogenous region highlighted in blue, haplotype specific methylation highlighted in red and the location of the structural variant indicated by the dashed line. **(Bottom left)** Panel showing methylation clustering of individual reads at the most hypomethylated region of the CDR revealing only a single cluster. **(Bottom right)** Panel showing methylation clustering of a more distal region of the CDR revealing haplotype specific methylation. We found three reads which spanned both the distal CDR region (red) and the SV (dotted line); one of these reads was hypermethylated and contained the SV, the other two were hypomethylated and did not have the deletion. **c)** The DXZ4 locus on CHM13 clustered into two haplotypes (presumably Xa and Xi), based solely on promoter methylation within each repeat unit.

Long-read sequencing also makes it possible to robustly assign reads to haplotypes, allowing us to generate a fully phased human epigenome. The longer the read, the greater the chance of encountering one or more heterozygous single-nucleotide polymorphisms (SNPs), which can be used to phase the reads into maternal or paternal origin (*17, 52, 53*). Because HG002 is a diploid cell line, we could evaluate allele-specific epigenetic states of HG002 across the T2T-CHM13 reference using variant data on HG002 and both parents(*54*). We were able to confidently phase 96.47% of the genome, including the haploid sex chromosomes. Variant calling in satellite repeats is unreliable, making phasing in these “SNP desert” regions difficult, however when reads are long enough they can be anchored to SNPs at the unique flanking regions. In HG002 we identified allele specific methylation throughout complex satellite arrays, such as the subtelomeric D4Z4 repeat associated with facioscapulohumeral dystrophy (*55*) **(Fig S17)**.

CHM13 contains nearly identical haplotypes which precludes SNP-based phasing techniques (*56*). However, using a methylation based clustering approach (Methods) allows us to identify haplotype specific methylation even in SNP deserts. Applying this analysis to an autosomal CDR, we examined the heterogeneity we previously noted in chromosome 19 (**Fig 3c**). We found only a single cluster at the center of the CDR, but identified two equally sized clusters in a region slightly distal to the CDR (26.32-26.35 Mb), suggesting the presence of haplotype specific methylation**(Fig 5b)**. We were able to anchor three of the clustered reads using a heterozygous structural variant ∼200 kb from the q-side edge of the CDR(*56*), validating these clusters as allele specific(**Fig 5b)**.

Looking outside the centromere, we applied single-read methylation analysis to the DXZ4 locus of the X chromosome, a 165 kb macrosatellite repeat containing 3 kb monomeric units (*57*). Each monomer of DXZ4 contains a bidirectional promoter with a CTCF site that is hypomethylated on the inactive X allele (Xi) and hypermethylated on the active X (Xa) (*58*). An initial analysis of the CHM13 sequencing reads in the region revealed distinct bands of hypermethylation and hypomethylation, with hypomethylation observed at the distal edge. Clustering reads by methylation at the promoter region of each DXZ4 monomer revealed two distinct clusters of methylated promoters and unmethylated promoters that we hypothesize correspond with the Xi allele and the Xa allele (**Fig 5c**). However, we observed some of the array’s promoters are hypomethylated across both clusters. We attribute this heterogeneity in array methylation to the early developmental state of CHM13 (*59*–*61*). For comparison, we looked at the Xa in HG002 and did not uncover any epigenetic heterogeneity within the array, perhaps due to the fact that it is a terminally differentiated male cell line **(Fig S18)**. Our results indicate that it is possible to interrogate allele-specific methylation events across large tandem repeats even in the absence of heterozygous variants. Clustering using the “fifth base” or 5mC has utility in regions or samples with low heterozygosity and allows for clear interrogation of haplotype specific behavior.

## DISCUSSION

Our analysis has provided a first glimpse into uncharted areas of the epigenome, leveraging the newly complete human genome assembly, T2T-CHM13. We found that this new genome, paired with dynamic k-mer assisted mapping methods, increased unique alignment of ENCODE ChIP-seq data by 2.35% in CHM13 compared to GRCh38. We reported increases in unique alignments across all histone marks surveyed and CTCF, indicating the increased reference quality of T2T-CHM13 and broadening the scope of existing epigenetic data to incorporate new genomic regions.

While short-read data coupled with k-mer assisted mapping can reveal gross enrichment in highly repetitive genomic regions, to perform detailed probing of specific epigenetic patterns we needed longread data. Leveraging our ability to measure methylation directly on nanopore long-reads, we interrogated the frequency of 5mC at 99.37% of the 32.3 million CpG sites within T2T-CHM13, 58.28% of which were contained within repetitive DNA, establishing the CHM13 methylome as the most complete human methylome surveyed to date (*62*). We compared methylation patterns within the highly complex pericentromeric HSat repeats between CHM13 and HG002 to a level of detail unobtainable with WGBS. We found that the tandemly repeated genomic sequences were reflected in the methylation patterns with clear and regular repeats in bands of hypo- and hyper-methylation. These bands may serve as functional genomic features (*i.e*. promoters and/or protein-binding domains). Pericentromeric satellites are likely to be rich in DNA-protein interactions to mediate secondary structure; they are known for their involvement in higher-order chromosome territory organization and the formation of the chromocenter (*63*–*65*). Alternatively, the methylation banding could be a result of sequence preference of *de novo* methyltransferases DNMT3A and DNMT3B (*66*) or could be influenced by DNA shape (*67, 68*). This result merits future studies into the functional role of methylation periodicity within pericentromeric satellites.

Within the HOR array, we identified a clearly demarcated hypomethylated region, the centromeric dip region (CDR), which is a universal feature of CHM13 centromeres. Though divergence in the HOR array precludes analysis of HG002 long-read data in the autosomes, using an assembly of the haploid X chromosome in HG002, we identified the same CDR feature in its centromere, establishing its presence in both of these cell lines. NanoNOMe chromatin accessibility data in HG002 reinforces the CDRs importance, showing extreme inaccessibility coordinated with the hypomethylated dip. We found that CENP-B motifs are largely unmethylated in the CDR and methylated outside the CDR, suggesting control of this element may be coordinated by the methylation. Finally, using marker-assisted mapping on CENP-A CUT&RUN data in both HG002 and CHM13, we observe localization of CENP-A to the CDR, further emphasizing its importance in kinetochore regulation.

One of the significant features of our single-molecule epigenetic data is our ability to investigate patterns of epigenetics; in the past, we and others have used this feature to phase contiguous blocks of the human epigenome (*17, 53*). We have taken this a step further, using methylation alone to cluster reads in repetitive areas devoid of heterozygous polymorphisms, including the DXZ4 array where the methylation signature is important to X chromosome inactivation (*55, 69*). We can also use this data to investigate the epigenetic heterogeneity within the centromere, observing heterogeneous patterns of methylation which may provide clues to how methylation is regulated at the CDR.

Our results highlight the power of long-read single-molecule assays to uncover novel epigenetic signatures. However, a significant challenge to further exploring the epigenome in a larger and more diverse sample set is optimal sequence alignment, especially among structurally variable HOR regions. Efforts by the Human Pangenome Reference Consortium to generate fully phased diploid genome assemblies within a few years will help to mitigate this problem. Clearly, we are currently only scratching the surface of what we can explore in the repetitive portions of the genome revealed by the new assembly. But many epigenetic assays still use short-read sequencing methods that, while useful with k-mer assisted mapping methods, still preclude clear measurement across repetitive regions or assessment of heterogeneity. Our work highlights the potential of long-read epigenetic methods to fully access the epigenetic landscape of repetitive regions of the genome. Following the examples set for chromatin accessibility(*17, 70*–*72*), other sequencing techniques, such as ChIP-seq and Hi-C can be adapted to utilize long reads revealing more complex patterns about genome regulation. A long-read multi-layered assay would not only provide simultaneous epigenetic data in repetitive DNA, but also further reveal information about the interaction of multiple epigenetic marks on single molecules.

## METHODS

### CHM13 T2T Nanopore

Human genomic DNA was extracted from the CHM13 cultured cell line. DNA extraction and sequencing methods are outlined in Miga et al, 2020; Logsdon et al., 2021 and Nurk et al. (*3, 45, 73*). Data is available at: **https://github.com/nanopore-wgs-consortium/CHM13**.

### CHM13 CpG Methylation Processing

CHM13 ultra long nanopore reads were aligned to the CHM13 reference with Winnowmap-v2.0 (*74*) with a k-mer size of 15. BAM files were filtered for primary alignments with SAMtools (v1.9) and filtered for read lengths greater than 50 kb. To measure CpG methylation in nanopore data we used Nanopolish (v0.13.2)(*16*). Nanopolish uses a Hidden Markov model on the nanopore current signal to distinguish 5mC from unmethylated cytosine. The methylation caller generates a log-likelihood value for the ratio of probability of methylated to unmethylated CGs at a specific k-mer. We next filtered methylation calls using the nanopore_methylation_utilities tool (https://github.com/timplab/nanopore-methylation-utilities), which uses a log-likelihood ratio of 1.5 as a threshold for calling methylation. CpG sites with log-likelihood ratios greater than 1.5 (methylated) or less than −1.5 (unmethylated) were considered high quality and included in the analysis. Reads that did not have any high-quality CpG sites were excluded from the subsequent methylation analysis. Nanopore_methylation_utilities integrates methylation information into the alignment BAM file for viewing in the bisulfite mode in Integrative Genomics Viewer (IGV) and also creates Bismark-style files(*75*). Methylation data was plotted by binning the genome with the BSgenome R package (BSgenome_1.56.0) and taking the average of CG sites within each bin. Single-read plots were generated with the ggplot2 R package (ggplot2_3.3.3) using the single-read data in the tabix indexed single-read methylation bed files generated from nanopore_methylation_utilities.

### HG002 Cell Culture

NA24385 cells (HG002) were obtained from the Coriell Institute (https://www.coriell.org/). Cells were grown in T-25 flasks in RPMI 1640 media with L-glutamine (Gibco; 11875093) supplemented with 15% fetal bovine serum (Gibco; 26140079) and 1% penicillin-streptomycin (Gibco; 15140122). Cells were cultured at 37C with 5% CO_2_ and were maintained by passaging ∼1/3 into fresh media every three days. Cells tested negative for mycoplasm contamination with the LookOut Mycoplasm PCR Detection Kit (Sigma; MP0035) and Jumpstart Taq DNA Polymerase (Sigma; D9307). Cells at passage 11 were used in nanoNOMe sequencing.

### NanoNOMe HG002 Sequencing

NanoNOMe library preparation was performed according to the methods outlined in Lee *et al*., 2020 (*17*). Cells were collected by resuspension, then nuclei were extracted by incubating in resuspension buffer (100 mM Tris-Cl, pH 7.4, 100 mM NaCl, 30 mM MgCl2) with 0.25% NP-40 for 5 min on ice. Intact nuclei were collected by centrifugation for 5 min at 500*g* at 4 °C. Nuclei were subjected to a methylation labeling reaction using a solution of 1x M.CviPI Reaction Buffer (NEB), 300 mM sucrose, 96 μM *S*-adenosylmethionine (NEB) and 200 U of M.CviPI (NEB) in 500 μl volume per 500,000 nuclei. The reaction mixture was incubated at 37C with shaking on a thermomixer at 1,000 RPM. for 15 min. *S*-adenosylmethionine was replenished at 96 μM at 7.5 min into the reaction. The reaction was stopped by the addition of an equal volume of stop solution (20 mM Tris-Cl, pH 7.9, 600 mM NaCl, 1% SDS, 10 mM disodium EDTA). Samples were treated with proteinase K (NEB) at 55C for >2 h and DNA was extracted via phenol:chloroform extraction and ethanol precipitation. After ethanol precipitation we enriched for HMW DNA with Circulomics Short Read Eliminator Extra Long (SRE-XL; SS-100-111-01).

Purified gDNA was prepared for nanopore sequencing following the protocol in the genomic sequencing by ligation kit LSK-SQK109 (ONT). Each nanopore library was prepared with 2 μg of input DNA. Fifteen libraries were generated and run on five PromethION flow cells (three libraries per flow cell). Flow cells were flushed at 24 hours and 48 hours with the Oxford Nanopore’s Flowcell Wash kit (EXP-WSH003) and reloaded with fresh library. Sequencing runs ran for a total of 72hrs and were simultaneously basecalled with Guppy 4.0.11. All nanoNOMe data can be accessed at on Sequence Read Archive with BioProject Accession number PRJNA725525.

### NanoNOMe Methylation Processing

HG002 nanoNOMe reads were aligned to a reference genome consisting of CHM13 chromosomes 1-22, HG002 chromosome X and GrCh38 chromosome Y with Winnowmap-v2.0 (*74*) with a k-mer size of 15. BAM files were filtered for primary alignments with SAM flag -F 256 and filtered for read lengths greater than 20 kb. To measure CpG and GpC methylation in nanopore data we used Nanopolish (v0.13.2) on the nanonome branch https://github.com/jts/nanopolish/tree/nanonome (*16*). We set an LLR threshold of -1/1 for GpC methylation calls and -1.5/1.5 for CpG methylation calls. Reads that did not have any high-quality sites were excluded from the subsequent methylation analysis. Nanopore_methylation_utilities integrates methylation information into the alignment BAM file for viewing in the bisulfite mode in IGV and also creates Bismark-style files.

In order to choose the optimal bin size for accessibility analysis, we used the intrinsic smoothness test previously explained by Sobecki *et al*. (*76*). 15 kb bins chosen by intrinsic smoothness test were then investigated for possible bias in CG and GC coverage. To visualize the methylation and accessibility patterns, the bins were z-normalized across each chromosome. Nucleosome footprints were determined by counting the number of consecutive unlabelled GpC sites, or “Inaccessible Runs”.

### Smoothed methylation Maps

CpG methylation frequency for methylation map plots was generated by calculating the fraction of methylated reads to total coverage within bins in CHM13 or HG002 with the BSGenome Bioconductor package (https://bioconductor.org/packages/BSgenome). Multiples of three bins were further smoothed with the “rollmean” function from the R package Zoo (https://cran.r-project.org/web/packages/zoo/index.html)(*77*). For comparing CpG coverage between CHM13 and HG002 CpG calls aligned to the T2T-CHM13 reference we flagged bins with greater than 90% of their methylation calls falling outside the coverage threshold (10X-100X) in the methylation maps to denote where calls are lower quality. Single-read plots were generated in the ggplot2 R package. Quantification panels above single-read plots were generated by taking the average methylation at each position in 1000 bins per 50 kb.

### Methylation clustering

Methylation clustering was done by selecting all reads spanning a specific locus and using the mclust (v5.4.7) R package with the “VII” model to cluster methylation calls across the locus(*78*). Within mclust we specified G as between 1 and 9 clusters. Positions with methylation calls that did not pass threshold to be called methylated or unmethylated were assigned a value of 0.5.

### Methylation Processing Code Availability

Code for all CHM13 and HG002 CpG methylation and GpC methylation available: https://github.com/timplab/T2T-Epigenetics

### SNP based phasing

HG002 variant calls were generated from ultra-long nanopore sequencing reads with PEPPER/Deep-Varient (https://github.com/kishwarshafin/pepper) (*54*). HG002 NanoNOMe sequencing reads were phased with whatshap (v1.0) (*79*) command: phase --ignore-read-groups.

### CENP-B motif annotation

CENP-B sites in HG002 chromosome X were identified with fuzznuc software tool from EMBOSS (*80*) searching for the CENP-B consensus sequence as follows:

~~~
fuzznuc --sequence HG002_chromsomeX.fasta --pattern NTTCGNNNNANNCGGGN -complement
~~~

### CUT&RUN Library Generation

CUT&RUN was carried out as in Thakur and Henikoff 2018 (*81*), with some variations. Frozen pellets of 1.6 M HG002 cells or 1.2 M CHM cells were thawed on ice and centrifuged at 500x*g* for 5 minutes at 4C. Cells were washed with cold PBS twice. For nuclear extraction, each cell pellet was resuspended in 500 uL of Nuclear Extraction Buffer (NEB, 20 mM HEPES pH 7.9, 10 mM KCl, 0.5mM Spermidine, 0.1% NP40, 20% glycerol, Roche Proteinase Inhibitor tablets) by pipetting gently, and incubated on ice for 5 minutes. Cells were centrifuged and washed with Washing Buffer (WB, 20mM HEPES pH 7.5, 150mM NaCl, 0.5mM Spermidine, 0.1% BSA, 0.05% NP40, Roche Proteinase Inhibitor tablets), blocked in WB containing BSA, and incubated in primary antibody for 2h at 4C under rotation. Primary antibodies used were: mouse CENP-A (Abcam, ab13939), rabbit CENP-B (Abcam, ab25734).

Cells were washed twice with WB and incubated with pAG-MNase (Cell Signaling) for 1h at 4C under rotation. For pAG-MNase digestion, samples were incubated on ice-water (0C) for 10 minutes, and CaCl_2_ was added to 2mM as final concentration. Samples were incubated for 30 minutes at 0C). To stop digestion, an equal volume of 2X STOP solution (200 mM NaCl, 20 mM EDTA, 4 mM EGTA, 0.1% NP40) was added. To recover low-salt fragments, samples were incubated for 1h at 4C under rotation, centrifuged at 500x*g* for 5 minutes and supernatant collected and labeled as low-salt fraction.. RNase A was added following incubation for 20 minutes at 37C. Samples were treated with Proteinase K for 1h at 65C (or overnight), and DNA extraction was carried out with MasterPure Complete DNA Isolation kit (Lucigen) as indicated by the manufacturer. Samples were analyzed by a Fragment Analyzer.

For library preparation, 1.5 pg of Spike-in Yeast DNA was added (obtained from the Henikoff lab) and NEBNext Ultra II End repair/A-tailing and Ligation kits were used as indicated by the manufacturer. DNA was cleaned using AMPure XP beads and the PCR reaction was carried out using NEBNext Ultra II Q5 master mix and NEBNext multiplex oligos for Illumina (12 cycles with annealing/extension for 15 seconds at 65C). Libraries were cleaned using AMPure XP beads as indicated by the manufacturer. Libraries were sequenced using NovaSeq 50PE sequencing.

### ENCODE Dynamic k-mer assisted mapping

We selected several ChIP-seq datasets generated as part of the ENCODE project (*18*) choosing ChIP-seq samples with at least 100 bp paired-end sequencing data and at least one matching input control (Methods). These criteria yielded 96 total sequencing libraries (**Supplemental Data 1**). All ENCODE-generated raw FASTQ files were downloaded from the ENCODE data portal (*82*). Prior to mapping, reads originating from a single library were combined. Reads were mapped with Bowtie2 (v2.4.1)(*83*) as paired-end with the arguments “--no-discordant --no-mixed --very-sensitive --no-unal --omit-sec-seq--xeq --reorder”. Alignments were filtered using SAMtools (v1.10) (*84*) using the arguments “-F 1804 -f 2 -q 2” to remove unmapped or single end mapped reads and those with a mapping quality score less than 2. PCR duplicates were identified and removed with the Picard tools “mark duplicates” command (v2.22.1, http://broadinstitute.github.io/picard) and the arguments “VALIDATION_STRINGEN-CY=LENIENT ASSUME_SORT_ORDER=queryname REMOVE_DUPLICATES = true”.

Alignments were then filtered for the presence of unique k-mers. Specifically, for each alignment, reference sequences aligned with template ends were compared to a database of k-mers unique in the whole genome. For each end of the paired-end sequencing reads, the k-mer length was determined by finding the largest multiple of 5 less than or equal to the aligned reference sequence length. Alignments were discarded if no unique kmers occurred in either end of the read. k-mer databases were generated using KMC3 (v3.1.1)(*85*). Alignments from replicates were then pooled. Bigwig genome tracks were created using deepTools bamCoverage (v3.4.3)(*86*) with a bin size of 1bp and default for all other parameters. Across all cell lines and marks, initial Bowtie2 alignments only yielded a 0.6% average increase in aligned reads to CHM13 versus GRCh38p13, however after intersecting the alignments with unique variable-length k-mers, the percentage of new alignments increased to 2.33% on average **(Table S1)**.

Mapping differences were broken down into five categories: satellites, LINEs/SINEs, segmental duplications, all other repetitive element types, and non-repetitive sequence. The segmental duplication regions were obtained from Vollger et al(*87*). The remaining repeat categories were obtained from Hoyt et al(*88*). LINE and SINE regions were extracted from the repeat masker annotation, merged so there were no overlapping elements, and intersected with the segmental duplication intervals to remove overlaps. The satellite regions were obtained the same way with the addition of intersecting them with both the segmental duplication and LINE/SINE intervals. The remaining repeat intervals were defined as all repeat masker annotations, merged and intersected with the previous three interval sets. The non-repetitive regions were defined as all intervals not covered by one of the above four tracks.

Peak calls were made using MACS2 (v2.2.7.1)(*89*) with default parameters and estimated genome sizes 3.03e9 and 2.79e9 for chm13v1 and GRCh38p13, respectively. GRCh38p13 peak calls were lifted over to chm13v1 using the UCSC liftOver utility, the chain file created by the T2T consortium (citation), and the parameter “-minMatch=0.2”. Peak intersections were determined using bedtools (v2.26.0)(*90*) counting each liftOver peak only once if any intersection occurred.

Full pipeline is available at https://github.com/msauria/T2T_Encode_Analysis.

### Reduced representation bisulfite sequencing (RRBS)

RRBS raw sequencing data from early human embryos from Guo *et al*.(*39*), was obtained from SRA (GEO accession: GSE49828) with fastq-dump (v2.8.0, http://ncbi.github.io/sra-tools/). RRBS reads were trimmed with TrimGalore (v0.6.6, https://github.com/FelixKrueger/TrimGalore) using the “--rrbs” and “--paired” parameters. The reads were aligned with Bismark (v0.22.2)(*75*) using the “bismark” command with the key parameters “-p --bam --bowtie” to the CHM13 reference genome prepared by the Bismark command “bismark_genome_preparation” with default parameters. Methylation data was extracted using the Bismark command “bismark_methylation_extractor” with the following parameters: “-p --comprehensive --merge_non_CpG --bedGraph --gzip --remove_spaces --cytosine_ report”. Methylation calls from technical replicates of the same biological replicate were combined by using the Bismark command “bismark2bedGraph”with the following parameters: “--buffer_size 20G--remove_spaces”. RRBS and CHM13 methylation data were then imported into R using the “read. bismark” command from the “bsseq” package (v1.24.4)(*91*) using only CHM13 reference CpGs with the following parameters “strandCollapse = TRUE, rmZeroCov = FALSE”. CpG loci were retained if they were covered by at least one read in 90% of the samples analyzed. Percent methylation was used to compare the samples by Euclidean distance with the R function “dist” with default parameters. Samples were clustered using the R function “hclust” using the “ward.D” method. The dendrogram (Fig S10) was plotted using “ggdendrogram” with default parameters. Full pipeline is available at https://github.com/timplab/pwh_projects/tree/master/chm13_rrbs.

### HG002 Bisulfite

TruSeq Bisulfite FASTQs were collected from the epiQC study (*92*). Paired-end FASTQs were aligned with Bismark (v0.22.2) with default parameters to a reference CHM13 autosomes (chromosomes 1-22), HG002 T2T chromosome X and GRCh38 chromosome Y (https://github.com/FelixKrueger/Bismark)(*75*) using the “bismark” command with the key parameters “-p --bam --bowtie” to the reference genome prepared by the Bismark command “bismark_genome_preparation” with default parameters. Methylation data was extracted using the Bismark command “bismark_methylation_extractor” with the following parameters: “-p --comprehensive --merge_non_CpG --bedGraph --gzip --remove_spaces--cytosine_report”. CpG methylation frequency for methylation map plot was generated by calculating the fraction of methylated reads to total coverage from the bismark CpG coverage bed file within bins in HG002 chromosome X with the BSGenome Bioconductor package (https://bioconductor.org/packages/BSgenome). Multiples of three bins were further smoothed with the “rollmean” function from the R package Zoo (https://cran.r-project.org/web/packages/zoo/index.html)(*77*).

### CUT&RUN marker-assisted mapping

Marker-assisted mapping of CUT&RUN data to the same genome (CHM13 to T2T-CHM13 or HG002 to CHM13 autosomes (chromosomes 1-22), HG002 T2T chromosome X and GRCh38 chromosome Y) was performed according to the methods outlined in Altemose *et al*., (*44*). In brief, 150 bp paired-end CUT&RUN libraries were mapped with bwa-mem (*93*) and filtered with SAMtools (*94*) for unique 51-mers as follows:

~~~
bwa mem -k 50 -c 1000000
overlapSelect -overlapBases=51
~~~

Unique 51-mers were generated with Meryl software (*95*). This method differs from dynamic k-mer assisted mapping in that there is a set k-mer size of k=51 and indels are not accounted for. More stringency can be placed on reads generated from the same genome as the reference as is the case with the CUT&RUN and not with the ENCODE datasets.

### Repeat-Masking

RepeatMasker annotations were generated in Hoyt *et al*.(*88*), and are available for both T2T-CHM13 and GRCh38 on the T2T UCSC genome assembly hub (http://t2t.gi.ucsc.edu/chm13/hub/hub.txt).

### Centromere annotations

Centromere region repeat annotations are described in Altemose *et al*.(*44*), and are available on the T2T UCSC genome assembly hub (http://t2t.gi.ucsc.edu/chm13/hub/hub.txt).

## ACKNOWLEDGMENTS

This study was supported by grants from the NIH R01HG009190 (W.T.), F32 GM134558 (G.A.L.), R24 DK106766-01A1 (M.C.S.), U24HG010263 (M.C.S.), 1R01HG011274-01 and 1U01HG010971 (K.H.M.)

This work was supported, in part, by the Intramural Research Program of the National Human Genome Research Institute, National Institutes of Health (S.K., A.R., and A.M.P.). This work was supported, in part, by the Damon Runyon Postdoctoral Fellowship and PEW Latin American Fellowship (G.V.C.) and a HHMI Hanna Gray Fellowship (N.A.). We acknowledge Rachael Workman and Samantha Sholes for reading and editing the manuscript, Isac Lee for engaging in discussion about epigenetics and chromatin state and Circulomics for their help in ultra high molecular weight DNA extraction.

## AUTHOR CONTRIBUTIONS

K.H.M and W.T. conceived the study. W.T., A.M.P., K.H.M., and A.G. coordinated the collaboration. K.H.M. and N.A. performed repeat characterization and satellite DNA assembly. A.G., M.E.G.S., and M.C.S performed ENCODE mapping and analysis. G.V.C. generated HG002 and CHM13 CUT&RUN libraries with assistance from N.A. A.R. and S.K. generated ONT mappings and CHM13 methylation calls. S.J.H and R.J.O. performed TE and non-centromeric repeat annotations. M.R.V., G.A.L. and E.E.E. generated SegDup annotations A.R., K.H.M., G.A.L., and S.J.H. performed marker-assisted mapping of CUT&RUN data. A.G., P.W.H, and R.R. performed HG002 cell culture, nanoNOMe sequencing, GpC and CpG methylation calls and accessibility analyses. A.G. and W.T. did nanopore methylation analysis on CHM13 and HG002. A.G., W.T. and K.H.M. developed figures. A.G. and W.T. drafted the manuscript. All authors provided critical feedback and read and approved the final manuscript.

## REFERENCES

1. E. S. Lander, L. M. Linton, B. Birren, C. Nusbaum, M. C. Zody, J. Baldwin, K. Devon, K. Dewar, M. Doyle, W. FitzHugh, R. Funke, D. Gage, K. Harris, A. Heaford, J. Howland, L. Kann, J. Lehoczky, R. LeVine, P. McEwan, K. McKernan, J. Meldrim, J. P. Mesirov, C. Miranda, W. Morris, J. Naylor, C. Raymond, M. Rosetti, R. Santos, A. Sheridan, C. Sougnez, Y. Stange-Thomann, N. Stojanovic, A. Subramanian, D. Wyman, J. Rogers, J. Sulston, R. Ainscough, S. Beck, D. Bentley, J. Burton, C. Clee, N. Carter, A. Coulson, R. Deadman, P. Deloukas, A. Dunham, I. Dunham, R. Durbin, L. French, D. Grafham, S. Gregory, T. Hubbard, S. Humphray, A. Hunt, M. Jones, C. Lloyd, A. McMurray, L. Matthews, S. Mercer, S. Milne, J. C. Mullikin, A. Mungall, R. Plumb, M. Ross, R. Shownkeen, S. Sims, R. H. Waterston, R. K. Wilson, L. W. Hillier, J. D. McPherson, M. A. Marra, E. R. Mardis, L. A. Fulton, A. T. Chinwalla, K. H. Pepin, W. R. Gish, S. L. Chissoe, M. C. Wendl, K. D. Delehaunty, T. L. Miner, A. Delehaunty, J. B. Kramer, L. L. Cook, R. S. Fulton, D. L. Johnson, P. J. Minx, S. W. Clifton, T. Hawkins, E. Branscomb, P. Predki, P. Richardson, S. Wenning, T. Slezak, N. Doggett, J. F. Cheng, A. Olsen, S. Lucas, C. Elkin, E. Uberbacher, M. Frazier, R. A. Gibbs, D. M. Muzny, S. E. Scherer, J. B. Bouck, E. J. Sodergren, K. C. Worley, C. M. Rives, J. H. Gorrell, M. L. Metzker, S. L. Naylor, R. S. Kucherlapati, D. L. Nelson, G. M. Weinstock, Y. Sakaki, A. Fujiyama, M. Hattori, T. Yada, A. Toyoda, T. Itoh, C. Kawagoe, H. Watanabe, Y. Totoki, T. Taylor, J. Weissenbach, R. Heilig, W. Saurin, F. Artiguenave, P. Brottier, T. Bruls, E. Pelletier, C. Robert, P. Wincker, D. R. Smith, L. Doucette-Stamm, M. Rubenfield, K. Weinstock, H. M. Lee, J. Dubois, A. Rosenthal, M. Platzer, G. Nyakatura, S. Taudien, A. Rump, H. Yang, J. Yu, J. Wang, G. Huang, J. Gu, L. Hood, L. Rowen, A. Madan, S. Qin, R. W. Davis, N. A. Federspiel, A. P. Abola, M. J. Proctor, R. M. Myers, J. Schmutz, M. Dickson, J. Grimwood, D. R. Cox, M. V. Olson, R. Kaul, C. Raymond, N. Shimizu, K. Kawasaki, S. Minoshima, G. A. Evans, M. Athanasiou, R. Schultz, B. A. Roe, F. Chen, H. Pan, J. Ramser, H. Lehrach, R. Reinhardt, W. R. McCombie, M. de la Bastide, N. Dedhia, H. Blöcker, K. Hornischer, G. Nordsiek, R. Agarwala, L. Aravind, J. A. Bailey, A. Bateman, S. Batzoglou, E. Birney, P. Bork, D. G. Brown, C. B. Burge, L. Cerutti, H. C. Chen, D. Church, M. Clamp, R. R. Copley, T. Doerks, S. R. Eddy, E. E. Eichler, T. S. Furey, J. Galagan, J. G. Gilbert, C. Harmon, Y. Hayashizaki, D. Haussler, H. Hermjakob, K. Hokamp, W. Jang, L. S. Johnson, T. A. Jones, S. Kasif, A. Kaspryzk, S. Kennedy, W. J. Kent, P. Kitts, E. V. Koonin, I. Korf, D. Kulp, D. Lancet, T. M. Lowe, A. McLysaght, T. Mikkelsen, J. V. Moran, N. Mulder, V. J. Pollara, C. P. Ponting, G. Schuler, J. Schultz, G. Slater, A. F. Smit, E. Stupka, J. Szustakowki, D. Thierry-Mieg, J. Thierry-Mieg, L. Wagner, J. Wallis, R. Wheeler, A. Williams, Y. I. Wolf, K. H. Wolfe, S. P. Yang, R. F. Yeh, F. Collins, M. S. Guyer, J. Peterson, A. Felsenfeld, K. A. Wetterstrand, A. Patrinos, M. J. Morgan, P. de Jong, J. J. Catanese, K. Osoegawa, H. Shizuya, S. Choi, Y. J. Chen, J. Szustakowki, International Human Genome Sequencing Consortium, Initial sequencing and analysis of the human genome. Nature. 409, 860–921 (2001).

2. J. C. Venter, M. D. Adams, E. W. Myers, P. W. Li, R. J. Mural, G. G. Sutton, H. O. Smith, M. Yandell, C. A. Evans, R. A. Holt, J. D. Gocayne, P. Amanatides, R. M. Ballew, D. H. Huson, J. R. Wortman, Q. Zhang, C. D. Kodira, X. H. Zheng, L. Chen, M. Skupski, G. Subramanian, P. D. Thomas, J. Zhang, G. L. Gabor Miklos, C. Nelson, S. Broder, A. G. Clark, J. Nadeau, V. A. McKusick, N. Zinder, A. J. Levine, R. J. Roberts, M. Simon, C. Slayman, M. Hunkapiller, R. Bolanos, A. Delcher, I. Dew, D. Fasulo, M. Flanigan, L. Florea, A. Halpern, S. Hannenhalli, S. Kravitz, S. Levy, C. Mobarry, K. Reinert, K. Remington, J. Abu-Threideh, E. Beasley, K. Biddick, V. Bonazzi, R. Brandon, M. Cargill, I. Chandramouliswaran, R. Charlab, K. Chaturvedi, Z. Deng, V. Di Francesco, P. Dunn, K. Eilbeck, C. Evangelista, A. E. Gabrielian, W. Gan, W. Ge, F. Gong, Z. Gu, P. Guan, T. J. Heiman, M. E. Higgins, R. R. Ji, Z. Ke, K. A. Ketchum, Z. Lai, Y. Lei, Z. Li, J. Li, Y. Liang, X. Lin, F. Lu, G. V. Merkulov, N. Milshina, H. M. Moore, A. K. Naik, V. A. Narayan, B. Neelam, D. Nusskern, D. B. Rusch, S. Salzberg, W. Shao, B. Shue, J. Sun, Z. Wang, A. Wang, X. Wang, J. Wang, M. Wei, R. Wides, C. Xiao, C. Yan, A. Yao, J. Ye, M. Zhan, W. Zhang, H. Zhang, Q. Zhao, L. Zheng, F. Zhong, W. Zhong, S. Zhu, S. Zhao, D. Gilbert, S. Baumhueter, G. Spier, C. Carter, A. Cravchik, T. Woodage, F. Ali, H. An, A. Awe, D. Baldwin, H. Baden, M. Barnstead, I. Barrow, K. Beeson, D. Busam, A. Carver, A. Center, M. L. Cheng, L. Curry, S. Danaher, L. Davenport, R. Desilets, S. Dietz, K. Dodson, L. Doup, S. Ferriera, N. Garg, A. Gluecksmann, B. Hart, J. Haynes, C. Haynes, C. Heiner, S. Hladun, D. Hostin, J. Houck, T. Howland, C. Ibegwam, J. Johnson, F. Kalush, L. Kline, S. Koduru, A. Love, F. Mann, D. May, S. McCawley, T. McIntosh, I. McMullen, M. Moy, L. Moy, B. Murphy, K. Nelson, C. Pfannkoch, E. Pratts, V. Puri, H. Qureshi, M. Reardon, R. Rodriguez, Y. H. Rogers, D. Romblad, B. Ruhfel, R. Scott, C. Sitter, M. Smallwood, E. Stewart, R. Strong, E. Suh, R. Thomas, N. N. Tint, S. Tse, C. Vech, G. Wang, J. Wetter, S. Williams, M. Williams, S. Windsor, E. Winn-Deen, K. Wolfe, J. Zaveri, K. Zaveri, J. F. Abril, R. Guigó, M. J. Campbell, K. V. Sjolander, B. Karlak, A. Kejariwal, H. Mi, B. Lazareva, T. Hatton, A. Narechania, K. Diemer, A. Muruganujan, N. Guo, S. Sato, V. Bafna, S. Istrail, R. Lippert, R. Schwartz, B. Walenz, S. Yooseph, D. Allen, A. Basu, J. Baxendale, L. Blick, M. Caminha, J. Carnes-Stine, P. Caulk, Y. H. Chiang, M. Coyne, C. Dahlke, A. Mays, M. Dombroski, M. Donnelly, D. Ely, S. Esparham, C. Fosler, H. Gire, S. Glanowski, K. Glasser, A. Glodek, M. Gorokhov, K. Graham, B. Gropman, M. Harris, J. Heil, S. Henderson, J. Hoover, D. Jennings, C. Jordan, J. Jordan, J. Kasha, L. Kagan, C. Kraft, A. Levitsky, M. Lewis, X. Liu, J. Lopez, D. Ma, W. Majoros, J. McDaniel, S. Murphy, M. Newman, T. Nguyen, N. Nguyen, M. Nodell, S. Pan, J. Peck, M. Peterson, W. Rowe, R. Sanders, J. Scott, M. Simpson, T. Smith, A. Sprague, T. Stockwell, R. Turner, E. Venter, M. Wang, M. Wen, D. Wu, M. Wu, A. Xia, A. Zandieh, X. Zhu, The sequence of the human genome. Science. 291, 1304–1351 (2001).

3. S. Nurk, S. Koren, A. Rhie, M. Rautianen, A. v. Bzikadze, A. Mikheenko, M. R. Vollger, N. Altemose, L. Uralsky, A. Gershman, S. Aganezov, S. J. Hoyt, M. Diekhans, G. A. Logsdon, M. Alonge, S. E. Antonarakis, M. Borchers, G. G. Bouffard, S. Y. Brooks, G. V. Galdas, H. Cheng, C.-S. Chin, W. Chow, G. de Lima Leonardo, M. Y. Dennis, P. C. Dishuck, R. Durbin, T. Dvorkina, I. T. Fiddes, G. Formenti, R. S. Fulton, A. Fungtammasan, E. Garrison, P. G. S. Grady, T. A. Graves-Lindsay, I. M. Hall, N. F. Hansen, G. A. Hartley, M. Haukness, K. Howe, M. W. Hunkapiller, C. Jain, M. Jain, E. D. Jarvis, P. Kerpedjiev, M. Kirsche, M. Kolmogorov, J. Korlach, M. Kremitzki, H. Li, V. V. Maduro, T. Marschall, A. M. McCartney, R. C. McCoy, D. E. Miller, J. C. Mullikin, E. W. Myers, B. Paten, P. Peluso, D. Porubsky, T. Potapova, E. I. Rogaev, J. A. Rosenfeld, S. L. Salzberg, V. A. Schneider, J. Sedlazeck Fritz, K. Shafin, C. J. Shew, A. Shumate, Y. Sims, D. C. Soto, I. Sović, A. Streets, B. A. Sullivan, F. Thibaud-Nissen, J. Torrance, J. Wagner, B. P. Walenz, Wood Jonathan M. D, C. Xiao, S. M. Yan, A. C. Young, U. Surti, I. A. Alexandrov, P. A. Pevzner, J. L. Gerton, R. J. O’Neill, W. Timp, J. M. Zook, M. C. Schatz, E. E. Eichler, K. H. Miga, A. M. Phillippy, The complete sequence of a human genome. bioRxiv (2021).

4. A. Janssen, S. U. Colmenares, G. H. Karpen, Heterochromatin: Guardian of the Genome. Annu. Rev. Cell Dev. Biol. 34, 265–288 (2018).

5. R. S. Gieni, G. K. T. Chan, M. J. Hendzel, Epigenetics regulate centromere formation and kinetochore function. J. Cell. Biochem. 104, 2027–2039 (2008).

6. N. C. Riddle, The role of epigenetics in maintaining genome stability. Biochem.. 39, 12–15 (2017).

7. A. J. Bannister, T. Kouzarides, Regulation of chromatin by histone modifications. Cell Res. 21, 381–395 (2011).

8. D. Nicetto, K. S. Zaret, Role of H3K9me3 heterochromatin in cell identity establishment and maintenance. Curr. Opin. Genet. Dev. 55, 1–10 (2019).

9. R. C. Allshire, G. H. Karpen, Epigenetic regulation of centromeric chromatin: old dogs, new tricks? Nat. Rev. Genet. 9, 923–937 (2008).

10. T. W. Depinet, J. L. Zackowski, W. C. Earnshaw, S. Kaffe, G. S. Sekhon, R. Stallard, B. A. Sullivan, G. H. Vance, D. L. Van Dyke, H. F. Willard, A. B. Zinn, S. Schwartz, Characterization of neo-centromeres in marker chromosomes lacking detectable alpha-satellite DNA. Hum. Mol. Genet. 6, 1195–1204 (1997).

11. W. Ren, H. Fan, S. A. Grimm, Y. Guo, J. J. Kim, J. Yin, L. Li, C. J. Petell, X.-F. Tan, Z.-M. Zhang, J. P. Coan, L. Gao, L. Cai, B. Detrick, B. Çetin, Q. Cui, B. D. Strahl, O. Gozani, Y. Wang, K. M. Miller, S. E. O’Leary, P. A. Wade, D. J. Patel, G. G. Wang, J. Song, Direct readout of heterochromatic H3K9me3 regulates DNMT1-mediated maintenance DNA methylation. Proc. Natl. Acad. Sci. U. S. A. 117, 18439–18447 (2020).

12. T. Pastinen, Genome-wide allele-specific analysis: insights into regulatory variation. Nat. Rev. Genet. 11, 533–538 (2010).

13. B. Carter, K. Zhao, The epigenetic basis of cellular heterogeneity. Nat. Rev. Genet. 22, 235–250 (2021).

14. G. A. Logsdon, M. R. Vollger, P. Hsieh, Y. Mao, M. A. Liskovykh, S. Koren, S. Nurk, L. Mercuri, P. C. Dishuck, A. Rhie, L. G. de Lima, D. Porubsky, A. V. Bzikadze, M. Kremitzki, T. A. Graves-Lindsay, C. Jain, K. Hoekzema, S. C. Murali, K. M. Munson, C. Baker, M. Sorensen, A. M. Lewis, U. Surti, J. L. Gerton, V. Larionov, M. Ventura, K. H. Miga, A. M. Phillippy, E. E. Eichler, The structure, function, and evolution of a complete human chromosome 8. Cold Spring Harbor Laboratory (2020), p. 2020.09.08.285395.

15. T. E. P. Consortium, The ENCODE Project Consortium, The ENCODE (ENCyclopedia Of DNA Elements) Project. Science. 306 (2004), pp. 636–640.

16. J. T. Simpson, R. E. Workman, P. C. Zuzarte, M. David, L. J. Dursi, W. Timp, Detecting DNA cytosine methylation using nanopore sequencing. Nat. Methods. 14, 407–410 (2017).

17. I. Lee, R. Razaghi, T. Gilpatrick, M. Molnar, A. Gershman, N. Sadowski, F. J. Sedlazeck, K. D. Hansen, J. T. Simpson, W. Timp, Simultaneous profiling of chromatin accessibility and methylation on human cell lines with nanopore sequencing. Nature Methods (2020), doi:10.1038/s41592-020-01000-7.

18. ENCODE Project Consortium, An integrated encyclopedia of DNA elements in the human genome. Nature. 489, 57–74 (2012).

19. Noma K, C. D. Allis, S. I. Grewal, Transitions in distinct histone H3 methylation patterns at the heterochromatin domain boundaries. Science. 293, 1150–1155 (2001).

20. F. L. Sun, M. H. Cuaycong, S. C. Elgin, Long-range nucleosome ordering is associated with gene silencing in Drosophila melanogaster pericentric heterochromatin. Mol. Cell. Biol. 21, 2867–2879 (2001).

21. A. C. Bell, A. G. West, G. Felsenfeld, Insulators and boundaries: versatile regulatory elements in the eukaryotic genome. Science. 291, 447–450 (2001).

22. N. D. Heintzman, R. K. Stuart, G. Hon, Y. Fu, C. W. Ching, R. D. Hawkins, L. O. Barrera, S. Van Calcar, C. Qu, K. A. Ching, W. Wang, Z. Weng, R. D. Green, G. E. Crawford, B. Ren, Distinct and predictive chromatin signatures of transcriptional promoters and enhancers in the human genome. Nat. Genet. 39, 311–318 (2007).

23. M. P. Creyghton, A. W. Cheng, G. G. Welstead, T. Kooistra, B. W. Carey, E. J. Steine, J. Hanna, M. Lodato, G. M. Frampton, P. A. Sharp, L. A. Boyer, R. A. Young, R. Jaenisch, Histone H3K27ac separates active from poised enhancers and predicts developmental state. Proc. Natl. Acad. Sci. U. S. A. 107, 21931–21936 (2010).

24. S. J. B. Holwerda, W. de Laat, CTCF: the protein, the binding partners, the binding sites and their chromatin loops. Philos. Trans. R. Soc. Lond. B Biol. Sci. 368, 20120369 (2013).

25. Z. Deng, Z. Wang, N. Stong, R. Plasschaert, A. Moczan, H.-S. Chen, S. Hu, P. Wikramasinghe, R. V. Davuluri, M. S. Bartolomei, H. Riethman, P. M. Lieberman, A role for CTCF and cohesin in subtelomere chromatin organization, TERRA transcription, and telomere end protection. EMBO J. 31, 4165–4178 (2012).

26. A. Ambrosini, S. Paul, S. Hu, H. Riethman, Human subtelomeric duplicon structure and organization. Genome Biol. 8, R151 (2007).

27. I. L. De Silanes, O. Grana, M. L. De Bonis, O. Dominguez, D. G. Pisano, M. A. Blasco, Identification of TERRA locus unveils a telomere protection role through association to nearly all chromosomes. Nat. Commun. 5, 1–13 (2014).

28. G. Le Berre, V. Hossard, J. F. Riou, Repression of TERRA expression by subtelomeric DNA methylation is dependent on NRF1 binding. International journal of (2019) (available at https://www.mdpi.com/1422-0067/20/11/2791).

29. N. Iglesias, S. Redon, V. Pfeiffer, M. Dees, J. Lingner, B. Luke, Subtelomeric repetitive elements determine TERRA regulation by Rap1/Rif and Rap1/Sir complexes in yeast. EMBO Rep. 12, 587–593 (2011).

30. M. Karimzadeh, C. Ernst, A. Kundaje, M. M. Hoffman, Umap and Bismap: quantifying genome and methylome mappability. Nucleic Acids Res. 46, e120 (2018).

31. L. L. Hall, M. Byron, D. M. Carone, T. W. Whitfield, G. P. Pouliot, A. Fischer, P. Jones, J. B. Lawrence, Demethylated HSATII DNA and HSATII RNA Foci Sequester PRC1 and MeCP2 into Cancer-Specific Nuclear Bodies. Cell Rep. 18, 2943–2956 (2017).

32. M. Ehrlich, N. E. Hopkins, G. Jiang, J. S. Dome, M. C. Yu, C. B. Woods, G. E. Tomlinson, M. Chintagumpala, M. Champagne, L. Dillerg, D. M. Parham, J. Sawyer, Satellite DNA hypomethylation in karyotyped Wilms tumors. Cancer Genet. Cytogenet. 141, 97–105 (2003).

33. Y. Delpu, T. McNamara, P. Griffin, S. Kaleem, S. Narayan, C. Schildkraut, K. Miga, M. Tahiliani, Chromosomal rearrangements at hypomethylated Satellite 2 sequences are associated with impaired replication efficiency and increased fork stalling. Cold Spring Harbor Laboratory (2019), p. 554410.

34. V. A. Schneider, T. Graves-Lindsay, K. Howe, N. Bouk, H.-C. Chen, P. A. Kitts, T. D. Murphy, K. D. Pruitt, F. Thibaud-Nissen, D. Albracht, R. S. Fulton, M. Kremitzki, V. Magrini, C. Markovic, S. McGrath, K. M. Steinberg, K. Auger, W. Chow, J. Collins, G. Harden, T. Hubbard, S. Pelan, J. T. Simpson, G. Threadgold, J. Torrance, J. M. Wood, L. Clarke, S. Koren, M. Boitano, P. Peluso, H. Li, C.-S. Chin, A. M. Phillippy, R. Durbin, R. K. Wilson, P. Flicek, E. E. Eichler, D. M. Church, Evaluation of GRCh38 and de novo haploid genome assemblies demonstrates the enduring quality of the reference assembly. Genome Res. 27, 849–864 (2017).

35. W. Timp, J. Comer, A. Aksimentiev, DNA base-calling from a nanopore using a Viterbi algorithm. Biophys. J. 102, L37–9 (2012).

36. S. L. Amarasinghe, S. Su, X. Dong, L. Zappia, M. E. Ritchie, Q. Gouil, Opportunities and challenges in long-read sequencing data analysis. Genome Biol. 21, 30 (2020).

37. T. H. Bestor, D. Bourc’his, Genetics and epigenetics of hydatidiform moles. Nat. Genet. 38 (2006), pp. 274–276.

38. N. Kato, A. Kamataki, H. Kurotaki, Methylation profiles of imprinted genes are distinct between mature ovarian teratoma, complete hydatidiform mole, and extragonadal mature teratoma. Mod. Pathol. 34, 502–507 (2021).

39. H. Guo, P. Zhu, L. Yan, R. Li, B. Hu, Y. Lian, J. Yan, X. Ren, S. Lin, J. Li, X. Jin, X. Shi, P. Liu, X. Wang, W. Wang, Y. Wei, X. Li, F. Guo, X. Wu, X. Fan, J. Yong, L. Wen, S. X. Xie, F. Tang, J. Qiao, The DNA methylation landscape of human early embryos. Nature. 511, 606–610 (2014).

40. K. H. Miga, Centromeric Satellite DNAs: Hidden Sequence Variation in the Human Population. Genes. 10 (2019), doi:10.3390/genes10050352.

41. N. Altemose, K. H. Miga, M. Maggioni, H. F. Willard, Genomic characterization of large heterochromatic gaps in the human genome assembly. PLoS Comput. Biol. 10, e1003628 (2014).

42. A. Vlahos, T. Mansell, R. Saffery, B. Novakovic, Human placental methylome in the interplay of adverse placental health, environmental exposure, and pregnancy outcome. PLoS Genet. 15, e1008236 (2019).

43. H. Okae, H. Chiba, H. Hiura, H. Hamada, A. Sato, T. Utsunomiya, H. Kikuchi, H. Yoshida, A. Tanaka, M. Suyama, T. Arima, Genome-wide analysis of DNA methylation dynamics during early human development. PLoS Genet. 10, e1004868 (2014).

44. N. Altemose, et al, Genetic and epigenetic maps of endogenous human centromeres. bioRxiv (to appear).

45. K. H. Miga, S. Koren, A. Rhie, M. R. Vollger, A. Gershman, A. Bzikadze, S. Brooks, E. Howe, D. Porubsky, G. A. Logsdon, V. A. Schneider, T. Potapova, J. Wood, W. Chow, J. Armstrong, J. Fredrickson, E. Pak, K. Tigyi, M. Kremitzki, C. Markovic, V. Maduro, A. Dutra, G. G. Bouffard, A. M. Chang, N. F. Hansen, F. Thibaud-Nissen, A. D. Schmitt, J.-M. Belton, S. Selvaraj, M. Y. Dennis, D. C. Soto, R. Sahasrabudhe, G. Kaya, J. Quick, N. J. Loman, N. Holmes, M. Loose, U. Surti, R. A. Risques, T. A. Graves Lindsay, R. Fulton, I. Hall, B. Paten, K. Howe, W. Timp, A. Young, J. C. Mullikin, P. A. Pevzner, J. L. Gerton, B. A. Sullivan, E. E. Eichler, A. M. Phillippy, Telomere-to-telomere assembly of a complete human X chromosome. Nature, 735928 (2020).

46. V. A. Shepelev, L. I. Uralsky, A. A. Alexandrov, Y. B. Yurov, E. I. Rogaev, I. A. Alexandrov, Annotation of suprachromosomal families reveals uncommon types of alpha satellite organization in pericentromeric regions of hg38 human genome assembly. Genomics Data. 5 (2015), pp. 139–146.

47. L. Andronov, K. Ouararhni, I. Stoll, B. P. Klaholz, A. Hamiche, CENP-A nucleosome clusters form rosette-like structures around HJURP during G1. Nat. Commun. 10, 4436 (2019).

48. M. M. Valdivia, K. Hamdouch, M. Ortiz, A. Astola, CENPA a genomic marker for centromere activity and human diseases. Curr. Genomics. 10, 326–335 (2009).

49. Y. Tanaka, H. Kurumizaka, S. Yokoyama, CpG methylation of the CENP-B box reduces human CENP-B binding. FEBS Journal. 272 (2004), pp. 282–289.

50. J.-H. Kim, T. Ebersole, N. Kouprina, V. N. Noskov, J.-I. Ohzeki, H. Masumoto, B. Mravinac, B. A. Sullivan, A. Pavlicek, S. Dovat, S. D. Pack, Y.-W. Kwon, P. T. Flanagan, D. Loukinov, V. Lobanenkov, V. Larionov, Human gamma-satellite DNA maintains open chromatin structure and protects a transgene from epigenetic silencing. Genome Research. 19 (2009), pp. 533–544.

51. K. Skvortsova, E. Masle-Farquhar, P.-L. Luu, J. Z. Song, W. Qu, E. Zotenko, C. M. Gould, Q. Du, T. J. Peters, Y. Colino-Sanguino, R. Pidsley, S. S. Nair, A. Khoury, G. C. Smith, L. A. Miosge, J. H. Reed, J. G. Kench, M. A. Rubin, L. Horvath, O. Bogdanovic, S. M. Lim, J. M. Polo, C. C. Goodnow, C. Stirzaker, S. J. Clark, DNA Hypermethylation Encroachment at CpG Island Borders in Cancer Is Predisposed by H3K4 Monomethylation Patterns. Cancer Cell. 35, 297–314.e8 (2019).

52. S. Gigante, Q. Gouil, A. Lucattini, A. Keniry, T. Beck, M. Tinning, L. Gordon, C. Woodruff, T. P. Speed, M. E. Blewitt, M. E. Ritchie, Using long-read sequencing to detect imprinted DNA methylation. Nucleic Acids Res. 47, e46 (2019).

53. V. Akbari, J.-M. Garant, K. O’Neill, P. Pandoh, R. Moore, M. A. Marra, M. Hirst, S. J. M. Jones, Megabase-scale methylation phasing using nanopore long reads and NanoMethPhase. Genome Biol. 22, 68 (2021).

54. S. Aganezov, et al, A complete human reference genome improves variant calling for population and clinical genomics. bioRxiv (to appear).

55. R. J. L. F. Lemmers, P. J. van der Vliet, J. P. Vreijling, D. Henderson, N. van der Stoep, N. Voermans, B. van Engelen, F. Baas, S. Sacconi, R. Tawil, S. M. van der Maarel, Cis D4Z4 repeat duplications associated with facioscapulohumeral muscular dystrophy type 2. Hum. Mol. Genet. 27, 3488–3497 (2018).

56. A. M. McCartney, et al, Chasing Perfection: Validation and Polishing Strategies for Telomere-to-Telomere Genome Assemblies. bioRxiv (to appear).

57. J. Giacalone, J. Friedes, U. Francke, A novel GC–rich human macrosatellite VNTR in Xq24 is differentially methylated on active and inactive X chromosomes. Nat. Genet. 1, 137–143 (1992).

58. B. P. Chadwick, DXZ4 chromatin adopts an opposing conformation to that of the surrounding chromosome and acquires a novel inactive X-specific role involving CTCF and antisense transcripts. Genome Research. 18 (2008), pp. 1259–1269.

59. I. M. van den Berg, J. S. E. Laven, M. Stevens, I. Jonkers, R.-J. Galjaard, J. Gribnau, J. H. van Doorninck, X chromosome inactivation is initiated in human preimplantation embryos. Am. J. Hum. Genet. 84, 771–779 (2009).

60. T. Tukiainen, A.-C. Villani, A. Yen, M. A. Rivas, J. L. Marshall, R. Satija, M. Aguirre, L. Gauthier, M. Fleharty, A. Kirby, B. B. Cummings, S. E. Castel, K. J. Karczewski, F. Aguet, A. Byrnes, GTEx Consortium, Laboratory, Data Analysis & Coordinating Center (LDACC)—Analysis Working Group, Statistical Methods groups—Analysis Working Group, Enhancing GTEx (eGTEx) groups, NIH Common Fund, NIH/NCI, NIH/NHGRI, NIH/NIMH, NIH/NIDA, Biospecimen Collection Source Site—NDRI, Biospecimen Collection Source Site—RPCI, Biospecimen Core Resource— VARI, Brain Bank Repository—University of Miami Brain Endowment Bank, Leidos Biomedi- cal—Project Management, ELSI Study, Genome Browser Data Integration & Visualization—EBI, Genome Browser Data Integration & Visualization—UCSC Genomics Institute, University of California Santa Cruz, T. Lappalainen, A. Regev, K. G. Ardlie, N. Hacohen, D. G. MacArthur, Landscape of X chromosome inactivation across human tissues. Nature. 550, 244–248 (2017).

61. J. C. Moreira de Mello, G. R. Fernandes, M. D. Vibranovski, L. V. Pereira, Early X chromosome inactivation during human preimplantation development revealed by single-cell RNA-sequenc- ing. Sci. Rep. 7, 10794 (2017).

62. Roadmap Epigenomics Consortium, A. Kundaje, W. Meuleman, J. Ernst, M. Bilenky, A. Yen, A. Heravi-Moussavi, P. Kheradpour, Z. Zhang, J. Wang, M. J. Ziller, V. Amin, J. W. Whitaker, M. D. Schultz, L. D. Ward, A. Sarkar, G. Quon, R. S. Sandstrom, M. L. Eaton, Y.-C. Wu, A. R. Pfenning, X. Wang, M. Claussnitzer, Y. Liu, C. Coarfa, R. A. Harris, N. Shoresh, C. B. Epstein, E. Gjoneska, D. Leung, W. Xie, R. D. Hawkins, R. Lister, C. Hong, P. Gascard, A. J. Mungall, R. Moore, E. Chuah, A. Tam, T. K. Canfield, R. S. Hansen, R. Kaul, P. J. Sabo, M. S. Bansal, A. Carles, J. R. Dixon, K.-H. Farh, S. Feizi, R. Karlic, A.-R. Kim, A. Kulkarni, D. Li, R. Lowdon, G. Elliott, T. R. Mercer, S. J. Neph, V. Onuchic, P. Polak, N. Rajagopal, P. Ray, R. C. Sallari, K. T. Siebenthall, N. A. Sinnott-Armstrong, M. Stevens, R. E. Thurman, J. Wu, B. Zhang, X. Zhou, A. E. Beaudet, L. A. Boyer, P. L. De Jager, P. J. Farnham, S. J. Fisher, D. Haussler, S. J. M. Jones, W. Li, M. A. Marra, M. T. McManus, S. Sunyaev, J. A. Thomson, T. D. Tlsty, L.-H. Tsai, W. Wang, R. A. Waterland, M. Q. Zhang, L. H. Chadwick, B. E. Bernstein, J. F. Costello, J. R. Ecker, M. Hirst, A. Meissner, A. Milosavljevic, B. Ren, J. A. Stamatoyannopoulos, T. Wang, M. Kellis, Integrative analysis of 111 reference human epigenomes. Nature. 518, 317–330 (2015).

63. M. Jagannathan, R. Cummings, Y. M. Yamashita, A conserved function for pericentromeric satellite DNA. Elife. 7 (2018), doi:10.7554/eLife.34122.

64. M. Jagannathan, Y. M. Yamashita, Function of Junk: Pericentromeric Satellite DNA in Chromosome Maintenance. Cold Spring Harb. Symp. Quant. Biol. 82, 319–327 (2017).

65. R. Mayer, A. Brero, J. von Hase, T. Schroeder, T. Cremer, S. Dietzel, Common themes and cell type specific variations of higher order chromatin arrangements in the mouse. BMC Cell Biol. 6, 44 (2005).

66. S.-Q. Mao, S. M. Cuesta, D. Tannahill, S. Balasubramanian, Genome-wide DNA Methylation Signatures Are Determined by DNMT3A/B Sequence Preferences. Biochemistry. 59, 2541–2550 (2020).

67. S. Rao, T.-P. Chiu, J. F. Kribelbauer, R. S. Mann, H. J. Bussemaker, R. Rohs, Systematic predic- tion of DNA shape changes due to CpG methylation explains epigenetic effects on protein–DNA binding. Epigenetics Chromatin. 11, 6 (2018).

68. A. Lazarovici, T. Zhou, A. Shafer, A. C. Dantas Machado, T. R. Riley, R. Sandstrom, P. J. Sabo, Y. Lu, R. Rohs, J. A. Stamatoyannopoulos, H. J. Bussemaker, Probing DNA shape and methylation state on a genomic scale with DNase I. Proc. Natl. Acad. Sci. U. S. A. 110, 6376–6381 (2013).

69. E. M. Darrow, M. H. Huntley, O. Dudchenko, E. K. Stamenova, N. C. Durand, Z. Sun, S.-C. Huang, A. L. Sanborn, I. Machol, M. Shamim, A. P. Seberg, E. S. Lander, B. P. Chadwick, E. L. Aiden, Deletion of DXZ4 on the human inactive X chromosome alters higher-order genome architecture. Proc. Natl. Acad. Sci. U. S. A. 113, E4504–12 (2016).

70. Y. Wang, A. Wang, Z. Liu, A. L. Thurman, L. S. Powers, M. Zou, Y. Zhao, A. Hefel, Y. Li, J. Zabner, K. F. Au, Single-molecule long-read sequencing reveals the chromatin basis of gene expression. Genome Res. 29, 1329–1342 (2019).

71. Z. Shipony, G. K. Marinov, M. P. Swaffer, N. A. Sinott-Armstrong, J. M. Skotheim, A. Kundaje, W. J. Greenleaf, Long-range single-molecule mapping of chromatin accessibility in eukaryotes,, doi:10.1101/504662.

72. A. B. Stergachis, B. M. Debo, E. Haugen, L. S. Churchman, J. A. Stamatoyannopoulos, Single-mol- ecule regulatory architectures captured by chromatin fiber sequencing. Science. 368, 1449–1454 (2020).

73. G. A. Logsdon, M. R. Vollger, P. Hsieh, Y. Mao, M. A. Liskovykh, S. Koren, S. Nurk, L. Mercuri, P. C. Dishuck, A. Rhie, L. G. de Lima, T. Dvorkina, D. Porubsky, W. T. Harvey, A. Mikheenko, A. V. Bzikadze, M. Kremitzki, T. A. Graves-Lindsay, C. Jain, K. Hoekzema, S. C. Murali, K. M. Munson, C. Baker, M. Sorensen, A. M. Lewis, U. Surti, J. L. Gerton, V. Larionov, M. Ventura, K. H. Miga, A. M. Phillippy, E. E. Eichler, The structure, function and evolution of a complete human chromo- some 8. Nature (2021), doi:10.1038/s41586-021-03420-7.

74. C. Jain, A. Rhie, N. Hansen, S. Koren, A. M. Phillippy, A long read mapping method for highly repetitive reference sequences. Cold Spring Harbor Laboratory (2020), p. 2020.11.01.363887.

75. F. Krueger, S. R. Andrews, Bismark: a flexible aligner and methylation caller for Bisulfite-Seq applications. Bioinformatics. 27, 1571–1572 (2011).

76. M. Sobecki, C. Souaid, J. Boulay, V. Guerineau, D. Noordermeer, L. Crabbe, MadID, a Versatile Approach to Map Protein-DNA Interactions, Highlights Telomere-Nuclear Envelope Contact Sites in Human Cells. Cell Rep. 25, 2891–2903.e5 (2018).

77. A. Zeileis, G. Grothendieck, zoo: S3 Infrastructure for Regular and Irregular Time Series. Journal of Statistical Software, Articles. 14, 1–27 (2005).

78. L. Scrucca, M. Fop, T. B. Murphy, A. E. Raftery, mclust 5: Clustering, Classification and Density Estimation Using Gaussian Finite Mixture Models. R J. 8, 289–317 (2016).

79. M. Martin, M. Patterson, S. Garg, S. Fischer, N. Pisanti, G. W. Klau, A. Schöenhuth, T. Marschall, WhatsHap: fast and accurate read-based phasing. bioRxiv (2016), p. 085050.

80. P. Rice, I. Longden, A. Bleasby, EMBOSS: the European Molecular Biology Open Software Suite. Trends Genet. 16, 276–277 (2000).

81. J. Thakur, S. Henikoff, Unexpected conformational variations of the human centromeric chromatin complex. Genes Dev. 32, 20–25 (2018).

82. C. A. Davis, B. C. Hitz, C. A. Sloan, E. T. Chan, J. M. Davidson, I. Gabdank, J. A. Hilton, K. Jain, U. K. Baymuradov, A. K. Narayanan, K. C. Onate, K. Graham, S. R. Miyasato, T. R. Dreszer, J. S. Strattan, O. Jolanki, F. Y. Tanaka, J. M. Cherry, The Encyclopedia of DNA elements (ENCODE): data portal update. Nucleic Acids Res. 46, D794–D801 (2018).

83. B. Langmead, S. L. Salzberg, Fast gapped-read alignment with Bowtie 2. Nat. Methods. 9, 357–359 (2012).

84. H. Li, B. Handsaker, A. Wysoker, T. Fennell, J. Ruan, N. Homer, G. Marth, G. Abecasis, R. Durbin, 1000 Genome Project Data Processing Subgroup, The Sequence Alignment/Map format and SAMtools. Bioinformatics. 25, 2078–2079 (2009).

85. M. Kokot, M. Dlugosz, S. Deorowicz, KMC 3: counting and manipulating k-mer statistics. Bioinformatics. 33, 2759–2761 (2017).

86. F. Ramírez, D. P. Ryan, B. Grüning, V. Bhardwaj, F. Kilpert, A. S. Richter, S. Heyne, F. Dündar, T. Manke, deepTools2: a next generation web server for deep-sequencing data analysis. Nucleic Acids Res. 44, W160–5 (2016).

87. Mitchell R. Vollger, Xavi Guitart, Philip C. Dishuck, Ludovica Mercuri, William T. Harvey, Ariel Gershman, Mark Diekhans, Arvis Sulovari, Katherine M. Munson, Alexandra M. Lewis, Kendra Hoekzema, David Porubsky, Ruiyang Li, Sergey Nurk, Sergey Koren, Karen H. Miga, Adam M. Phillippy, Winston Timp, Mario Ventura, Evan E. Eichler, Segmental duplications and their variation in a complete human genome. bioRxiv (2021).

88. S. J. Hoyt, et al, From telomere to telomere: characterizing the transcriptional and epigenetic state of repeat elements. bioRxiv (to appear).

89. Y. Zhang, T. Liu, C. A. Meyer, J. Eeckhoute, D. S. Johnson, B. E. Bernstein, C. Nusbaum, R. M. Myers, M. Brown, W. Li, X. S. Liu, Model-based analysis of ChIP-Seq (MACS). Genome Biol. 9, R137 (2008).

90. A. R. Quinlan, I. M. Hall, BEDTools: a flexible suite of utilities for comparing genomic features. Bioinformatics. 26, 841–842 (2010).

91. K. D. Hansen, B. Langmead, R. A. Irizarry, BSmooth: from whole genome bisulfite sequencing reads to differentially methylated regions. Genome Biol. 13, R83 (2012).

92. J. Foox, J. Nordlund, C. Lalancette, T. Gong, M. Lacey, S. Lent, B. W. Langhorst, V. K. C. Ponnaluri, L. Williams, K. Padmamabhan, R. Cavalcante, A. Lundmark, D. Butler, J. Gurvitch, J. M. Greally, M. Suzuki, M. Menor, M. Nasu, A. Alonso, C. Sheridan, A. Scherer, S. Bruinsma, G. Golda, Muszynska, P. P. Labaj, M. A. Campbe, F. Wos, A. Raine, U. Liljedahl, T. Axelsson, C. Wang, Z. Chen, Z. Yang, J. Li, X. Yang, H. Wang, A. Melnick, S. Guo, A. Blume, V. Franke, I. Ibanez de Caceres, C. Rodriguez-Antolin, R. Rosas, J. Wade Davis, J. Ishii, D. B. Megherbi, W. Xiao, W. Liao, J. Xu, H. Hong, B. Ning, W. Tong, A. Akalin, Y. Wang, C. E. Mason, The SEQC2 Epigenomics Quality Control (EpiQC) Study: comprehensive characterization of epigenetic methods, reproducibility, and quantification. bioRxiv (2020),, doi:10.1101/2020.12.14.421529.

93. H. Li, R. Durbin, Fast and accurate short read alignment with Burrows-Wheeler transform. Bioinformatics. 25 (2009), pp. 1754–1760.

94. P. Danecek, J. K. Bonfield, J. Liddle, J. Marshall, V. Ohan, M. O. Pollard, A. Whitwham, T. Keane, S. A. McCarthy, R. M. Davies, H. Li, Twelve years of SAMtools and BCFtools. Gigascience. 10 (2021), doi:10.1093/gigascience/giab008.

95. J. R. Miller, A. L. Delcher, S. Koren, E. Venter, B. P. Walenz, A. Brownley, J. Johnson, K. Li, C. Mobarry, G. Sutton, Aggressive assembly of pyrosequencing reads with mates. Bioinformatics. 24, 2818–2824 (2008).

