## Supplemental Material for "Epigenetic Patterns in a Complete Human Genome"

**TABLE OF CONTENTS**

**SUPPLEMENTAL FIGURES:**

**Figure S1:** Overview of ENCODE dynamic k-mer mapping pipeline.
**Figure S2:** ENCODE mapping summary.
**Figure S3.** Dynamic k-mer mapping read filtering.
**Figure S4:** Enrichment of histone marks and CTCF across ENCODE cell lines.
**Figure S5:** Percent divergence of the Telomere Associated Repeat (TAR).
**Figure S6:** epiQC TruSeq Bisulfite data aligned to HG002 centromere X.
**Figure S7:** Difference in number of CpG sites identified between T2T-CHM13 and GRCh38.
**Figure S8:** Thresholds for long-read nanopore methylation.
**Figure S9:** CHM13 CpG methylation quality control.
**Figure S10:** Comparison of methylation in early human embryo samples and CHM13.
**Figure S11:** CpG call coverage thresholding.
**Figure S12:** Methylation frequency panel of CHM13 and HG002 data aligned to centromeric autosomes.
**Figure S13:** Repeat type CpG densities.
**Figure S14:** Methylation frequency of CHM13 centromeric regions.
**Figure S15:** NanoNOMe alignment and GpC methylation calling.
**Figure S16.** NanoNOMe Inaccessible Run Lengths.
**Figure S17:** Phased D4Z4 methylation frequency and accessibility in HG002.
**Figure S18:** DXZ4 methylation frequency in HG002.

**SUPPLEMENTAL TABLES:**

**Table S1:** Statistics of dynamic k-mer assisted mapping of ENCODE data.
**Table S2:** New CpG Regions in CHM13.
**Table S3.** Summary table of methylation high quality calls.
**Table S4.** Accessions of RRBS data.
**Table S5.** CpG site coverage thresholding.
**Table S6.** CHM13 repeat median methylation values.
**Table S7.** Median methylation in HG002 and CHM13 repeats on the X chromosome.

**SUPPLEMENTAL DATA:**

**Supplemental Data 1:** Summary of ENCODE data and results.

**Supplementary Figures:**

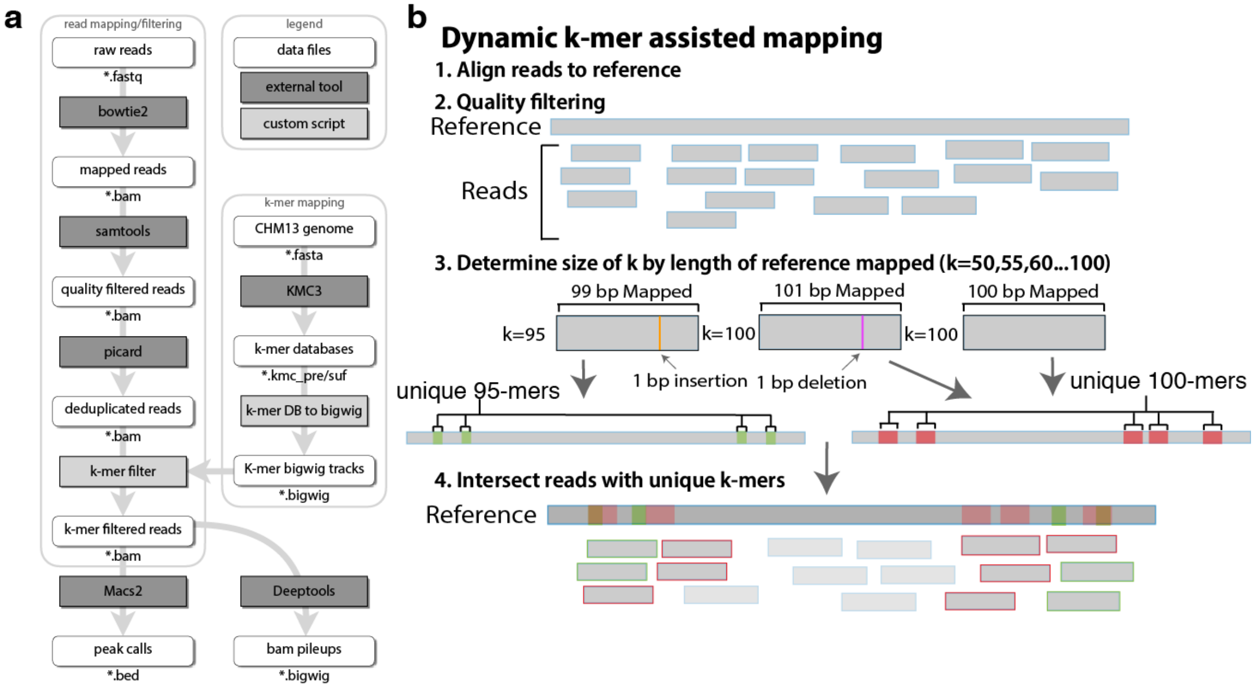


***Figure S1. Overview of ENCODE dynamic k-mer mapping pipeline. a)*** *Bioinformatic pipeline for mapping ENCODE data to repetitive regions of the T2T-CHM13 genome.* ***b)*** *Schematic description of the dynamic k-mer assisted mapping  pipeline. The size of the k-mers in the k-mer filtering step are dependent on the length of the mapped reference sequence. We generated k-mer databases for 50-100mers by multiples of five.  If all 100bp of a read map then the 100mer database is used. However, if the reference sequence is longer or shorter than 100bp then we use the database that is shorter than the reference length to the nearest five. For example, when there is a 1bp insertion in the read compared to reference, the corresponding reference sequence is 99bp, therefore the 95-mer database is used. When there is a 1bp deletion in the read compared to the reference, the corresponding reference sequence is 101 bp long so we use the 100-mer database. Mismatches only impact the k-mer size if they occur in the first or last positions, otherwise reference sequence length is unchanged.*

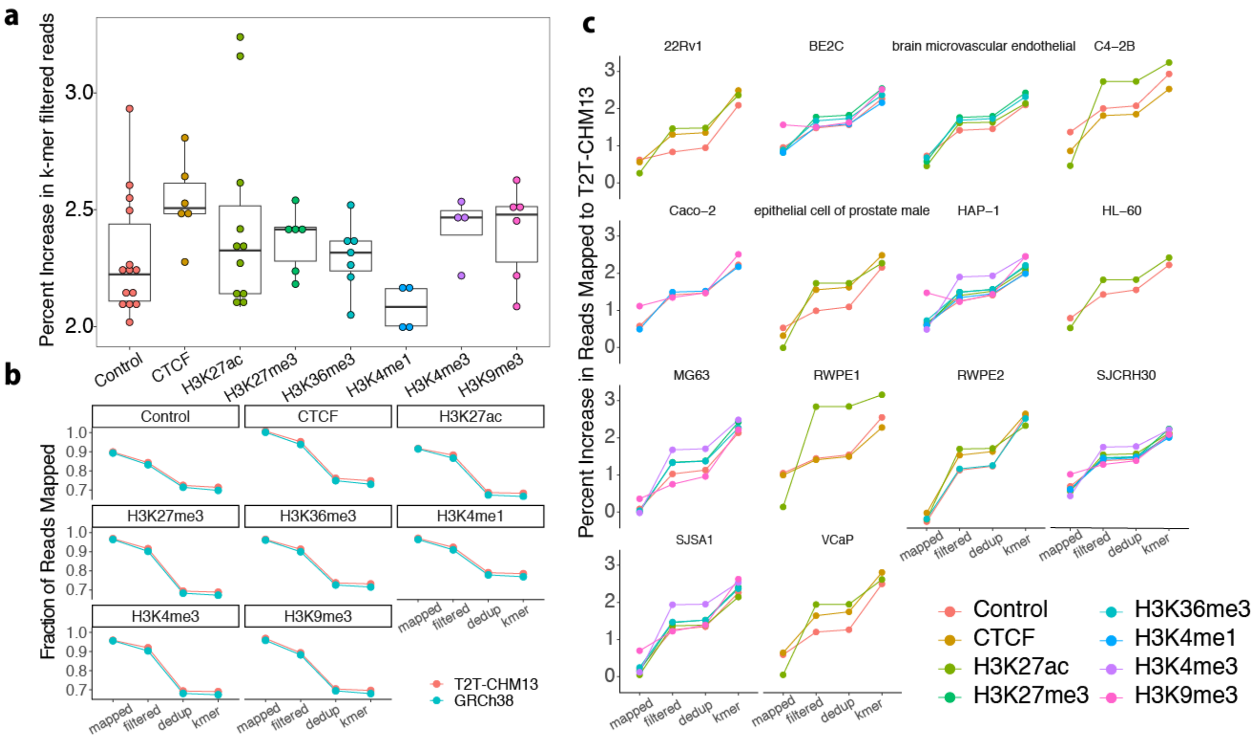


***Figure S2. ENCODE mapping summary. a)*** *Relative percent increase in k-mer filtered aligned reads in T2T-CHM13 compared to GRCh38 for all datasets surveyed. Each point represents an ENCODE cell line.* ***b)*** *Fraction of total aligned reads retained during each mapping step in the pipeline.* ***c)*** *Percent increase in alignments to T2T-CHM13 compared to GRCh38 at each mapping step separated by cell line.*

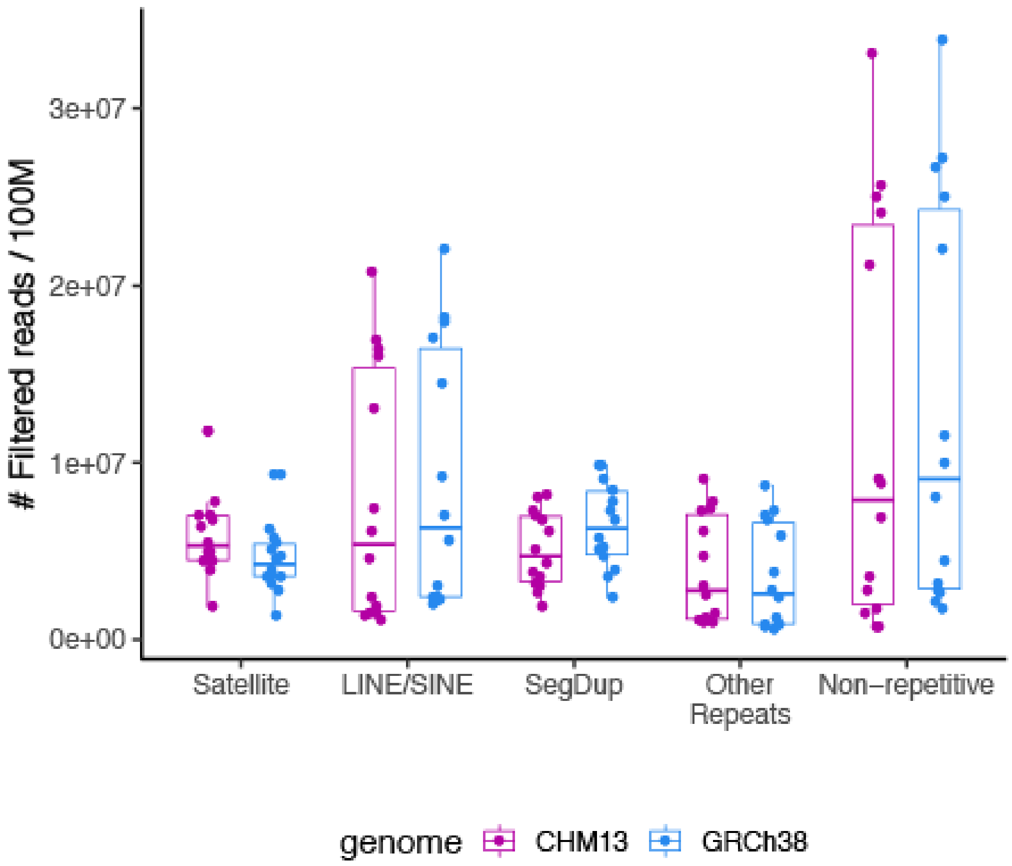


***Figure S3. Dynamic k-mer mapping read filtering.*** *Number of mapped reads filtered out of input control ChIP-seq library normalized to a post alignment read depth of 100M reads after quality filtering, deduplication and k-mer filtering in different genomic regions for each ENCODE cell line profiled.*

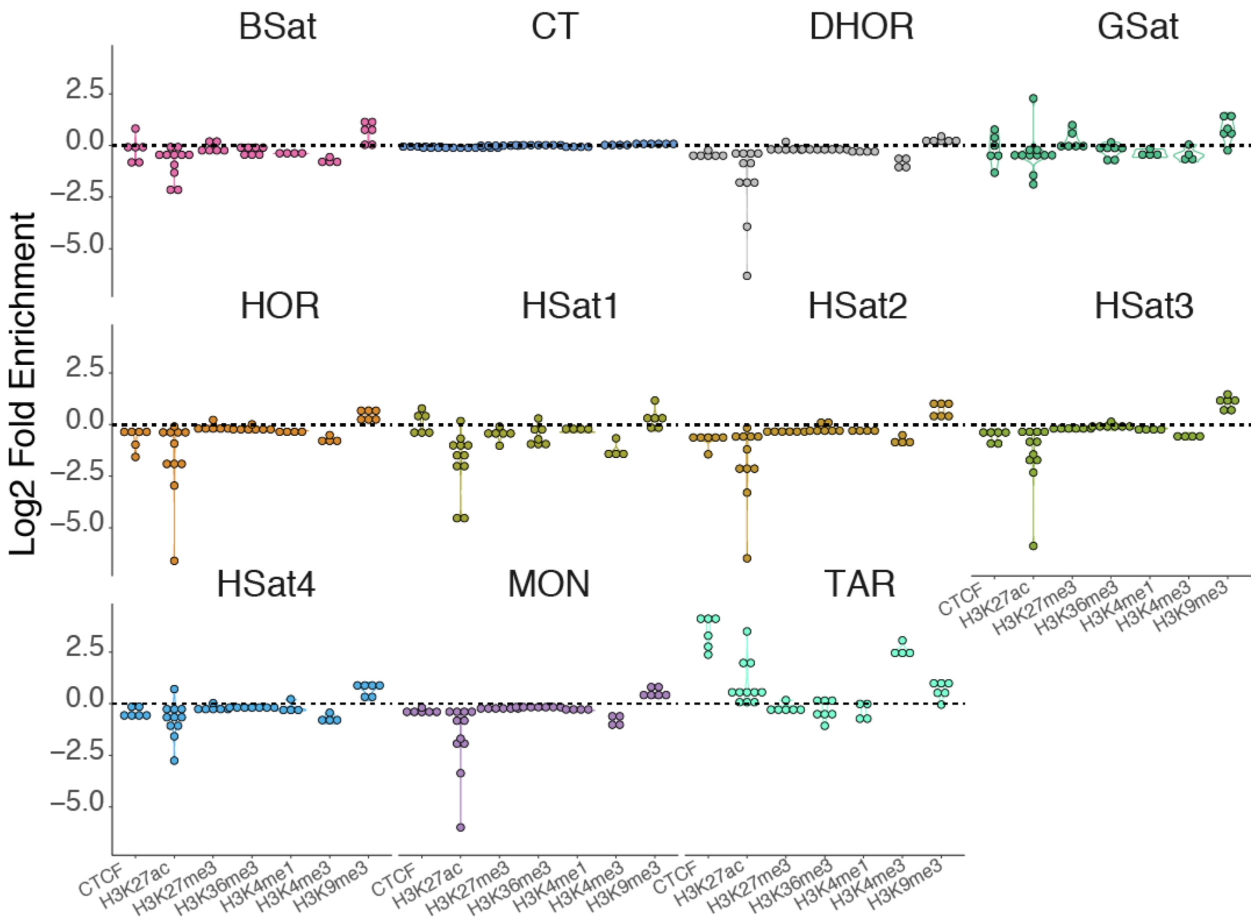


***Figure S4. Enrichment of histone marks and CTCF across ENCODE cell lines.*** *Log2 fold enrichment of epigenetic mark versus input control for each satellite repeat class.*

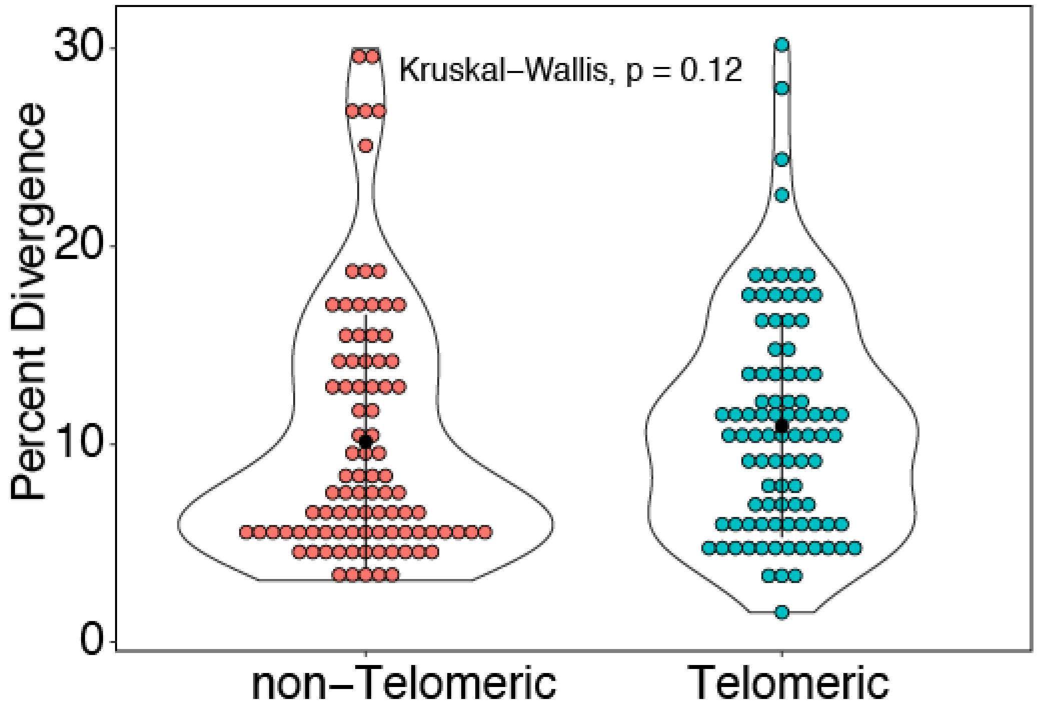


***Figure S5. Percent divergence of the Telomere Associated Repeat (TAR).*** *Telomeric TAR repeats are defined as within 2 kb of the chromosome end. TAR repeats outside this region are defined as non-telomeric. Percent divergence calculated by RepeatMasker from the TAR consensus sequence.*

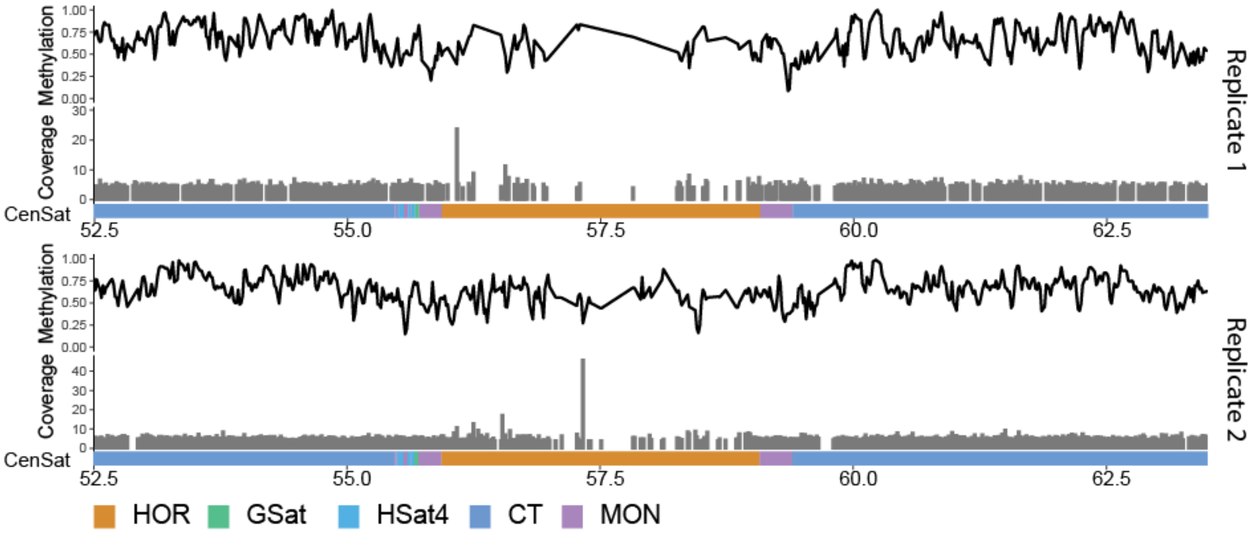


***Figure S6. epiQC TruSeq Bisulfite data aligned to HG002 centromere X.*** *Bismark alignments and methylation calls of the epiQC TruSeq Bisulfite data on HG002 to the HG002 chromosome X centromere (1). Coverage track is the number of times a CpG site is called, within the HOR region there is significant dropout of coverage.*

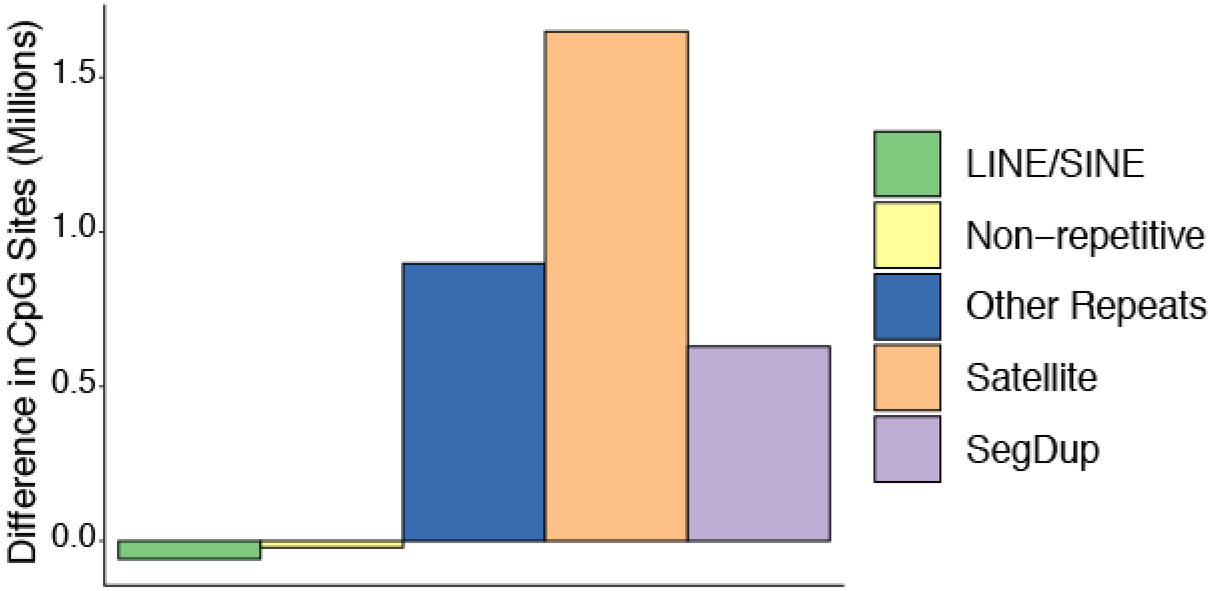


***Figure S7. Difference in number of CpG sites identified between T2T-CHM13 and GRCh38.***
*Positive value indicates more CpGs in T2T-CHM13, negative indicates more CpGs in GRCh38. Overall, GRCh38 has 29.2M CpGs (omitting chrY) versus T2T-CHM13's 32.3M CpGs.*

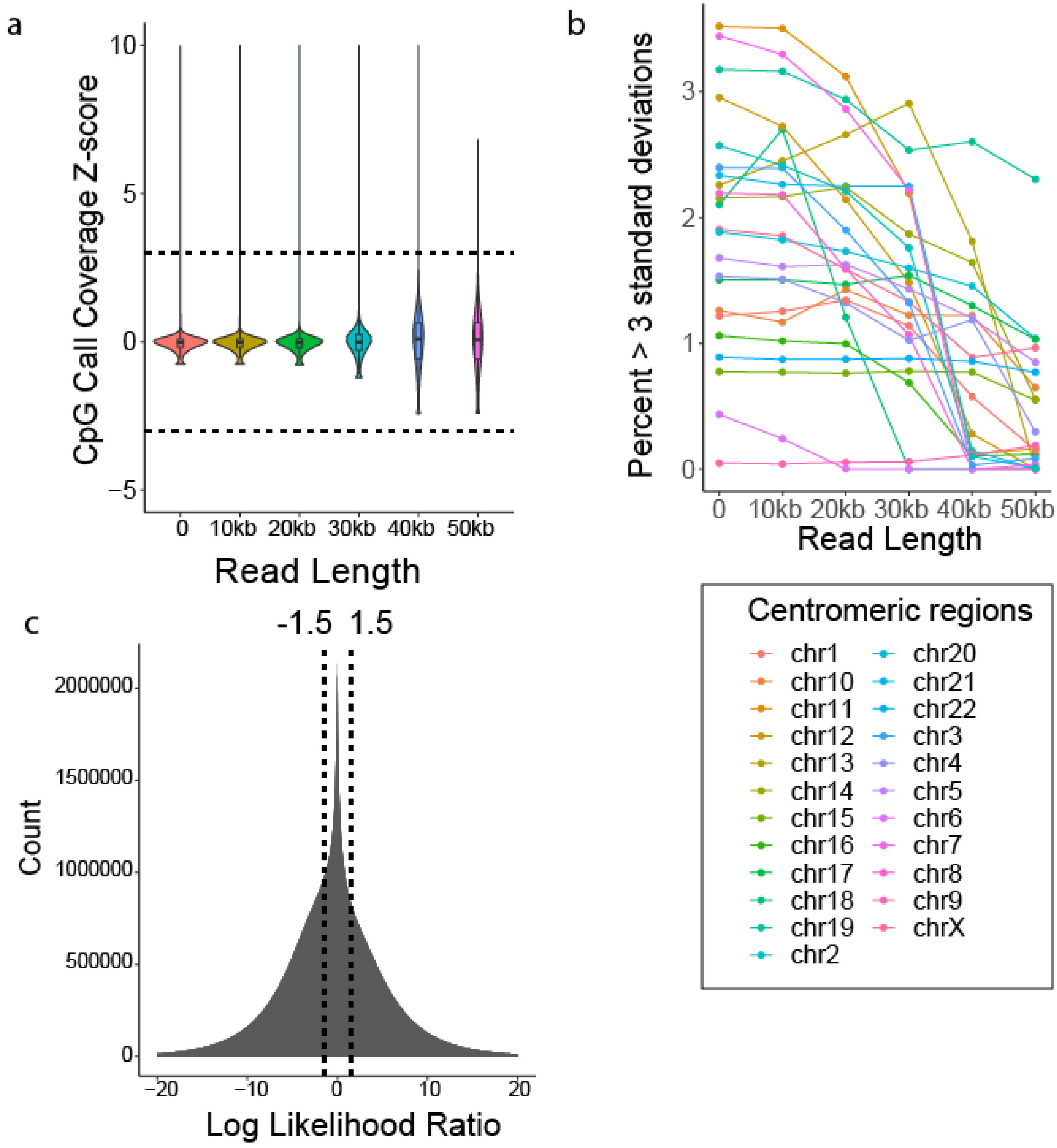


***Figure S8. Thresholds for long-read nanopore methylation. a-b)*** *Z-score of CpG coverage as compared against the whole genome. With increasing read length the percentage of CpGs with coverage Z-scores greater than 3 and less than -3 decreased. 50kb reads were chosen to decrease coverage bias in centromeric regions, allowing for robust analysis of methylation calls through centromeric repeats.* ***c)*** *The distribution of all log-likelihood scores for methylation calls in CHM13. Calls with log likelihood ratios greater than 1.5 and less than -1.5 were considered high quality and used for all subsequent analysis.*

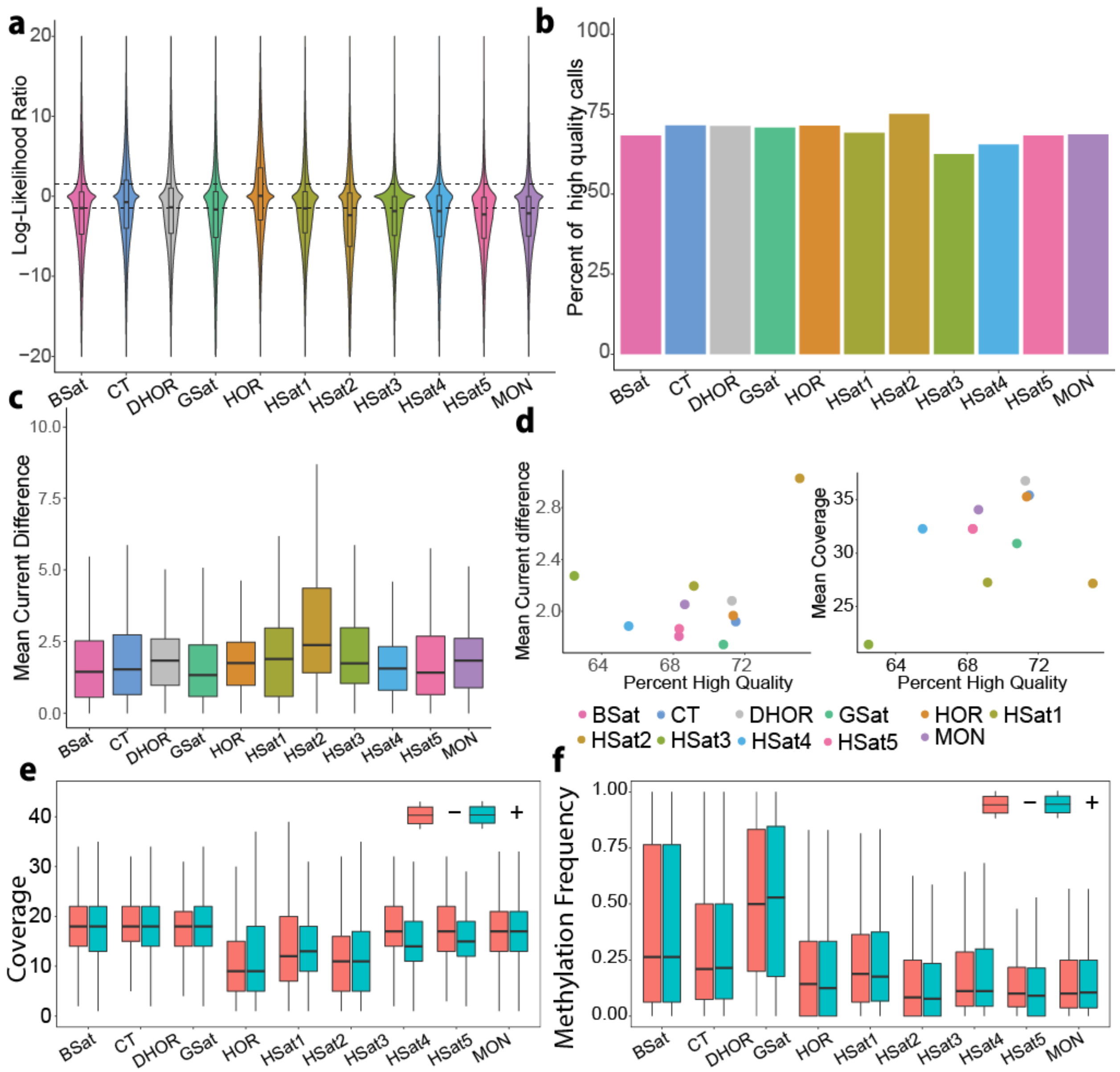


**Figure S9.** ***CHM13 CpG methylation quality control.*** ***a)*** *Distribution of log-likelihood ratios for each repeat type from nanopore reads filtered to only primary alignments >50 Kb.* ***b)*** *Percentage of high quality ( |log likelihood| > 1.5) CpG calls within each satellite repeat.* ***c)*** *Distribution of the mean current difference between methylated and unmethylated k-mers possible in each repeat. For all CG containing 6-mers per repeat the absolute value of the mean difference in methylated vs unmethylated current distribution was calculated. Boxplots are weighted by kmer frequency.* ***d)*** *(Left) Scatter plot of the percentage of high quality calls versus the mean current difference of methylated vs unmethylated 6-mers per repeat type weighted by kmer frequency, (Pearson's Correlation, R=.36, p=.28). (Right) Scatter plot of the percentage of high quality methylation calls versus CpG coverage per repeat, (Pearson's Correlation, R=.42, p=.19).* ***e)*** *Called CpG site coverage per read strand within each repeat type.* ***f)*** *Average CpG methylation frequency per read strand within each repeat type.*

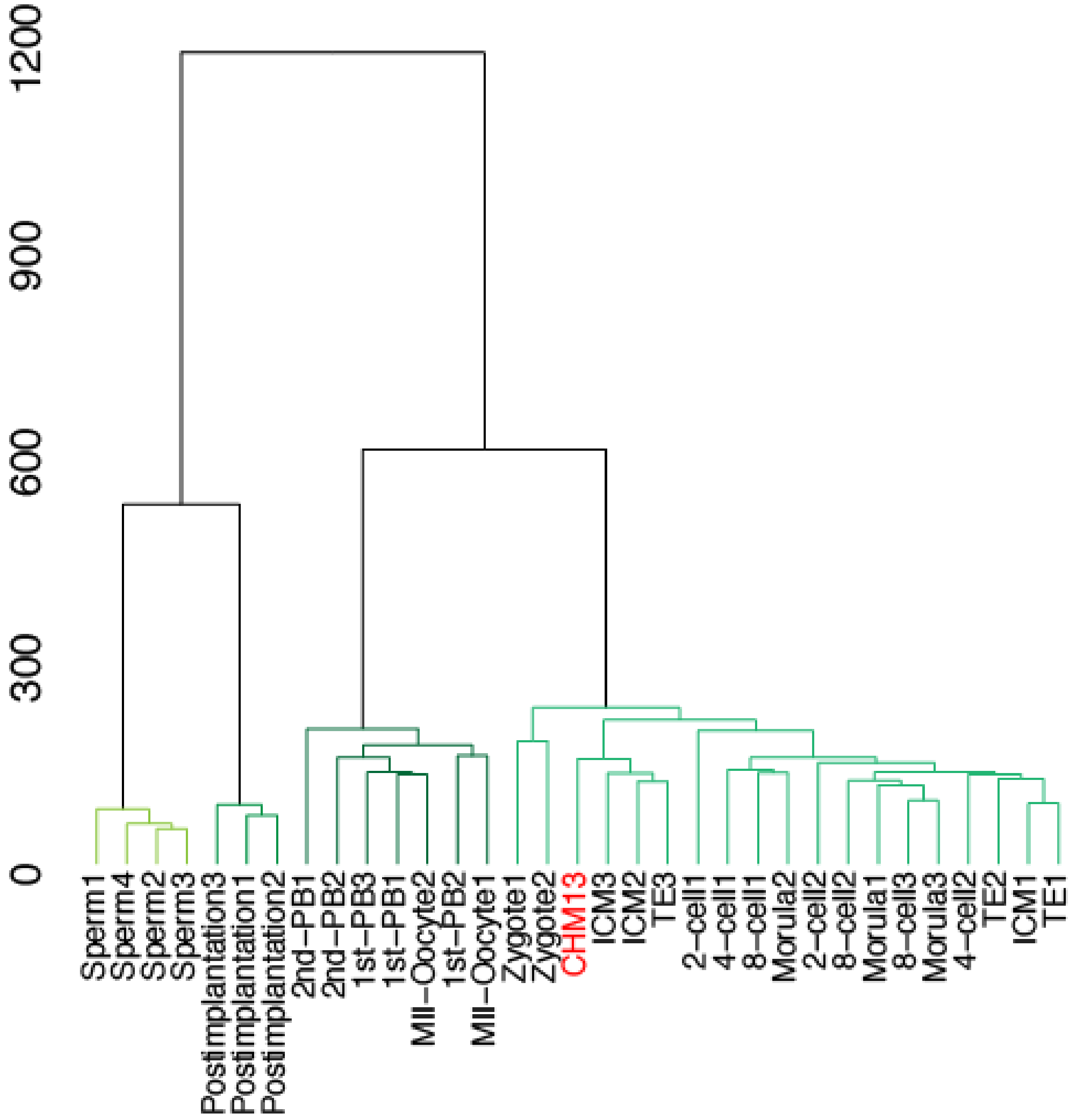


***Figure S10. Comparison of methylation in early human embryo samples and CHM13.*** *Reduced representation bisulfite sequencing (RRBS) methylation data from 12 stages of human embryo development(2) compared to CHM13 methylation generated from nanopore data. T2T-CHM13 reference CpGs covered by at least 1 read in 90% of samples were used. Raw methylation percentages were clustered using Euclidean distance and ward.D clustering then plotted as a dendrogram.*

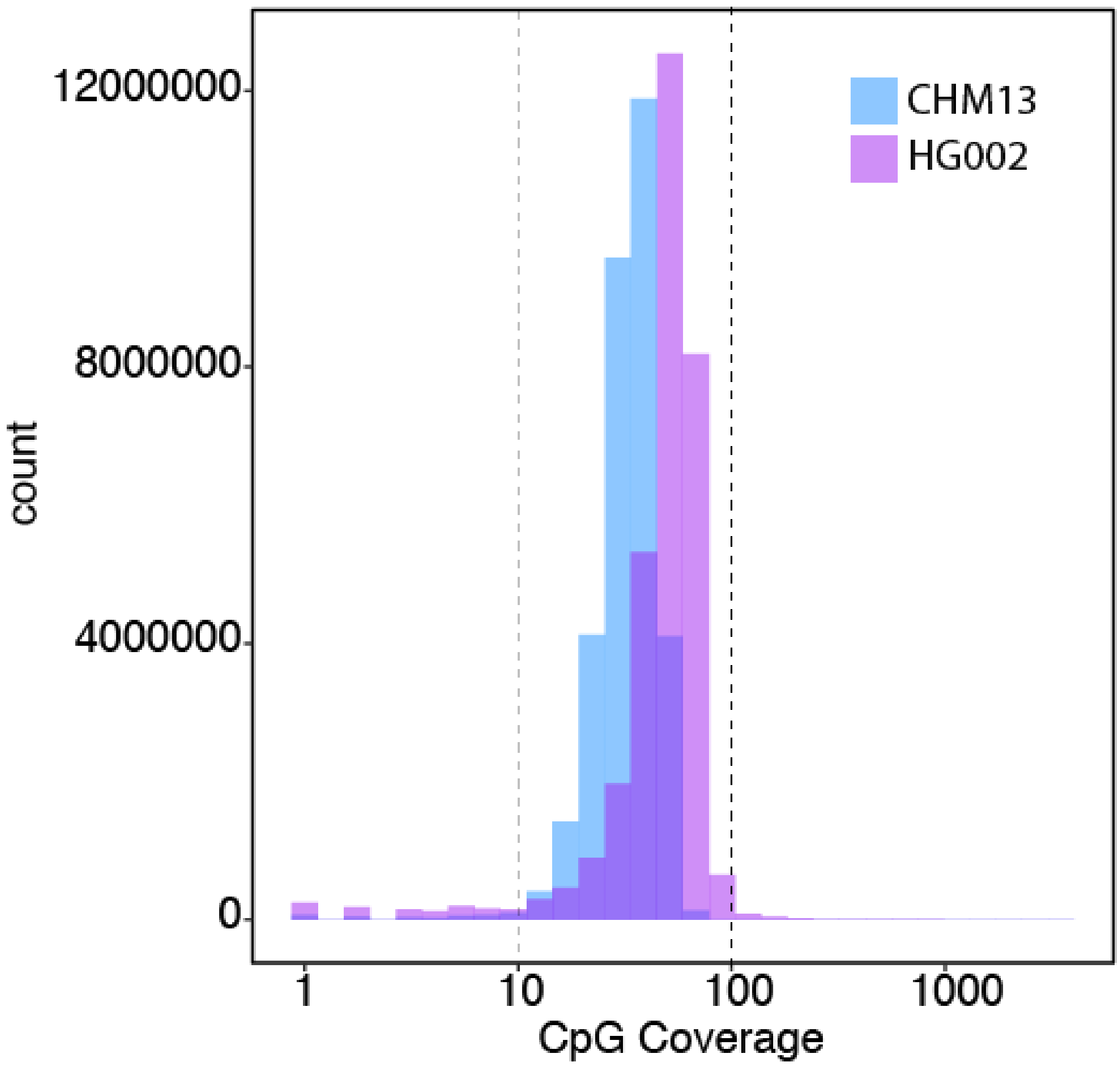


***Figure S11. CpG call coverage thresholding***. *Coverage histogram of number of methylation calls per CpG site. CHM13 data includes all reads longer than 50 Kb with high quality methylation calls. HG002 NanoNOMe data includes all reads longer than 20 Kb with high quality methylation calls. CpGs with coverage metrics between the dashed lines (10-100X) were considered high quality sites. CpG sites with coverage lower than 10X or higher than 100X were flagged as lower quality sites.*

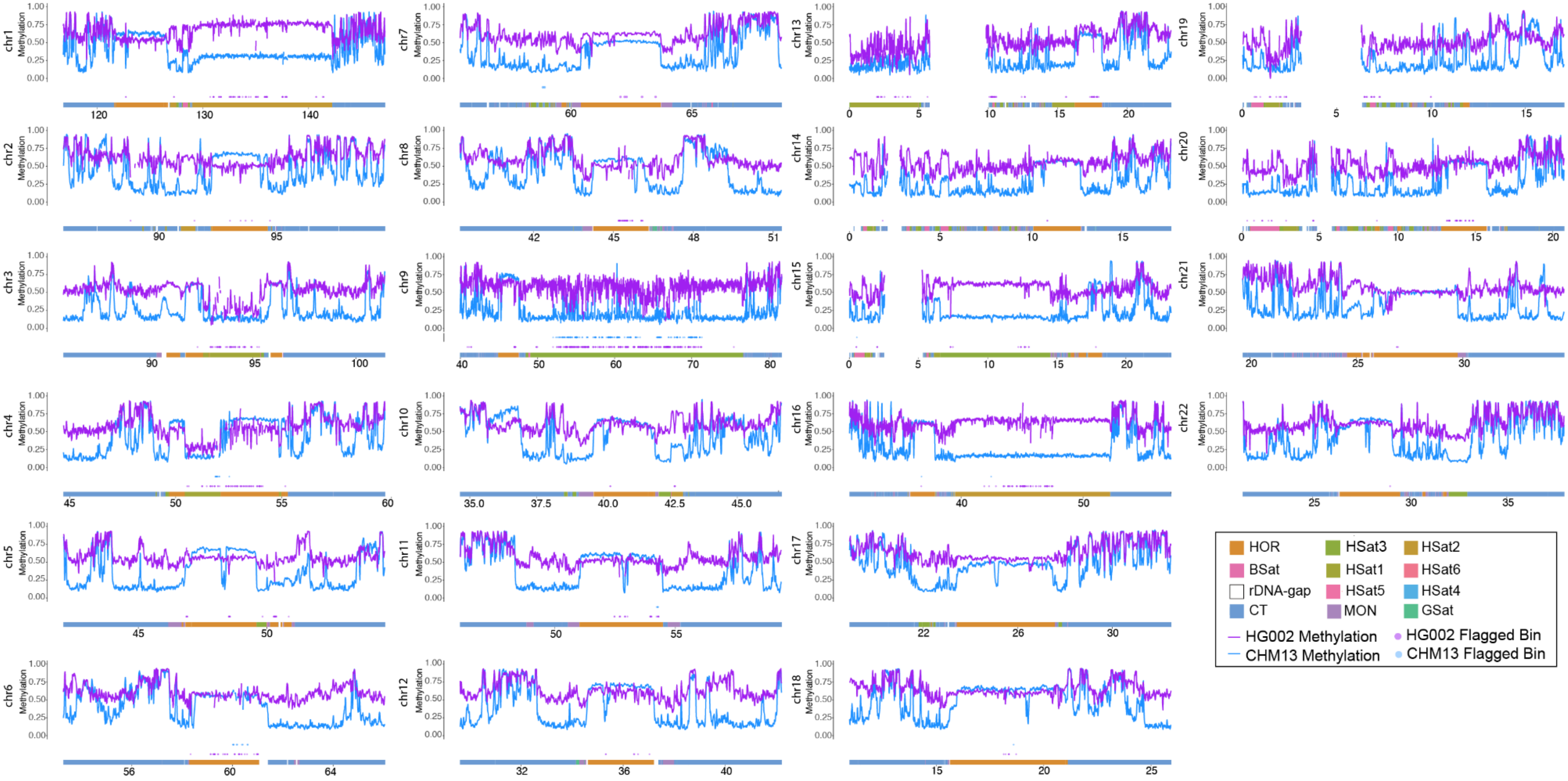


***Figure S12. Methylation frequency panel of CHM13 and HG002 data aligned to centromeric autosomes.*** *Methylation track represents mean methylation frequency in 10 Kb bins smoothed with a rolling average over three bins. Lower chromosome ideogram is colored with satellite repeat annotations. Points represent flagged 10 Kb bins with greater than 90% of their methylation calls outside the coverage threshold (10X-100X).*

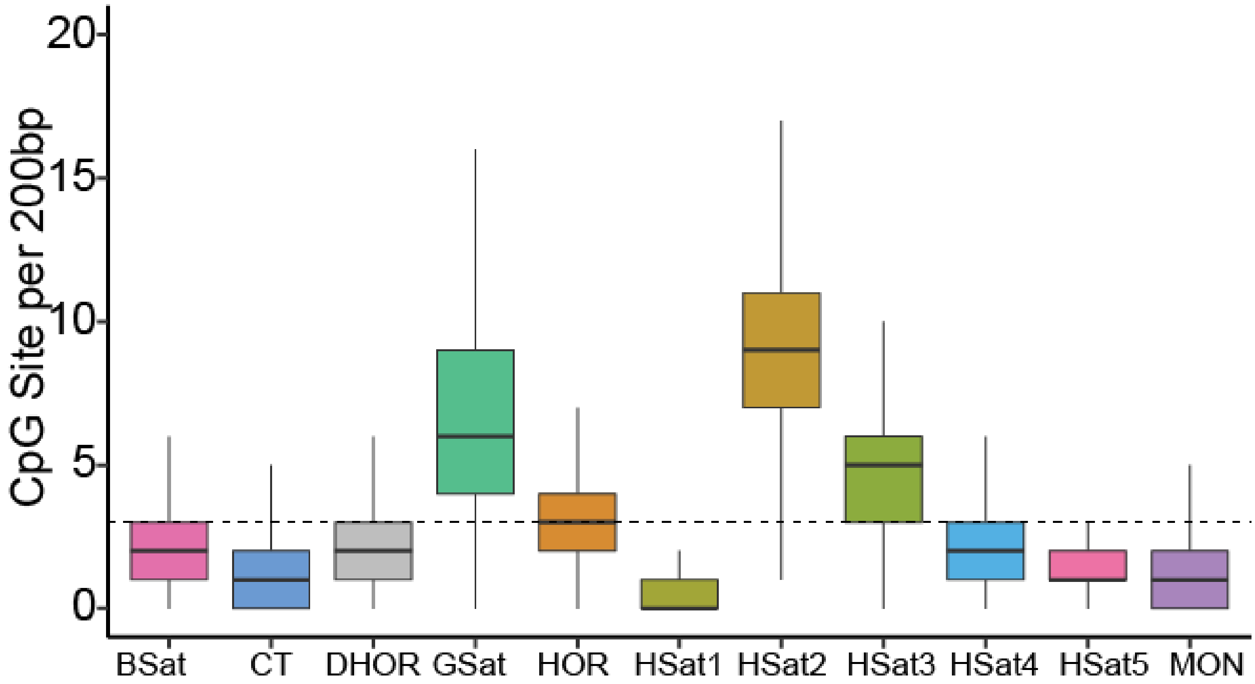


***Figure S13. Repeat type CpG densities.*** *CpG density measured per satellite repeat type using a 200bp sliding window with 1bp step size. Boxplots show the distribution of sliding window CpG sites. Dotted line represents the whole genome median of CpG sites per 200bp sliding window.*

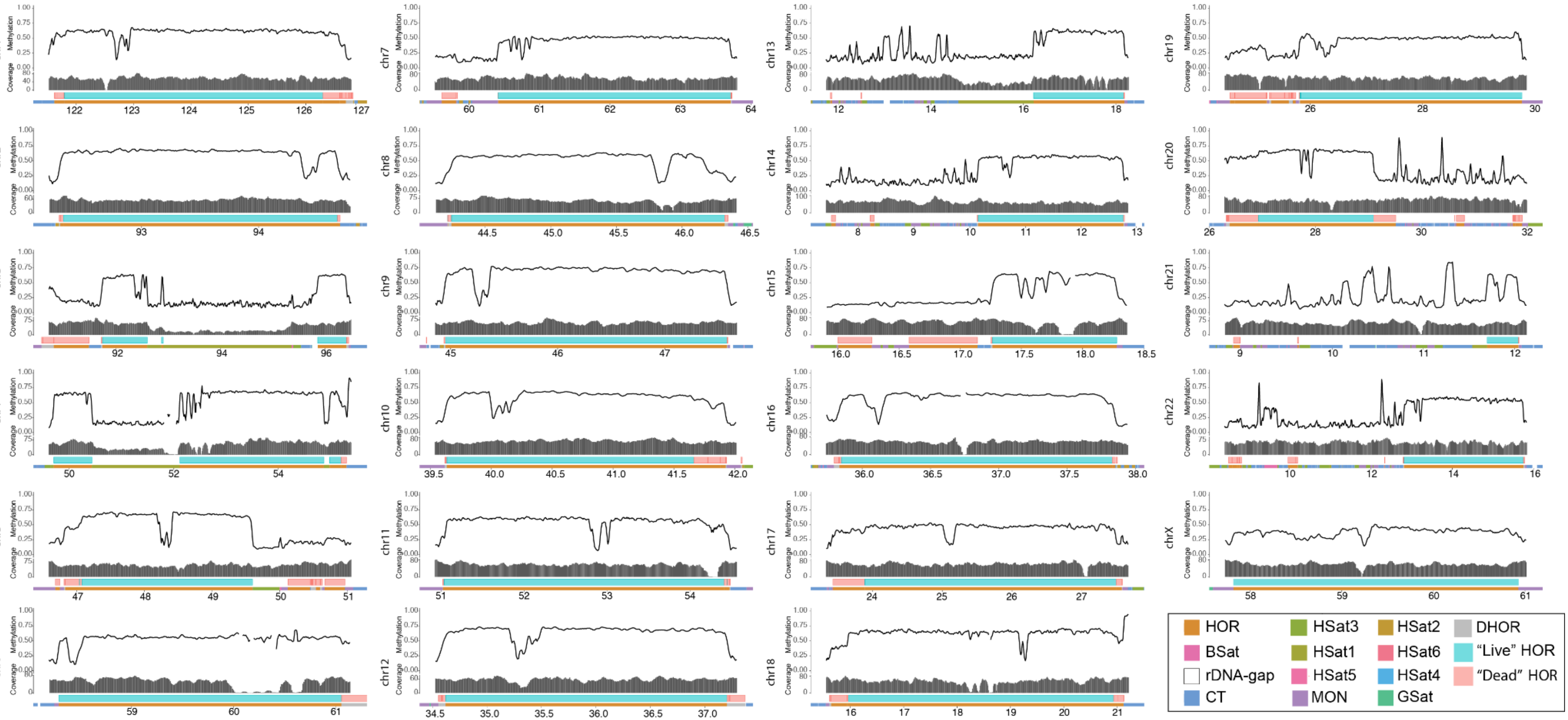


***Figure S14. Methylation frequency of CHM13 centromeric regions***. *Panels representing the centromeric regions of each chromosome of CHM13. CHM13 methylation frequency is plotted in 10 Kb bins smoothed with a rolling average over three bins. The depth panel represents the number of aligned nanopore reads containing at least one high quality CpG methylation call. Pink and gray boxes annotate the “live” and “dead” HOR arrays. Bottom line represents satellite repeat annotation*

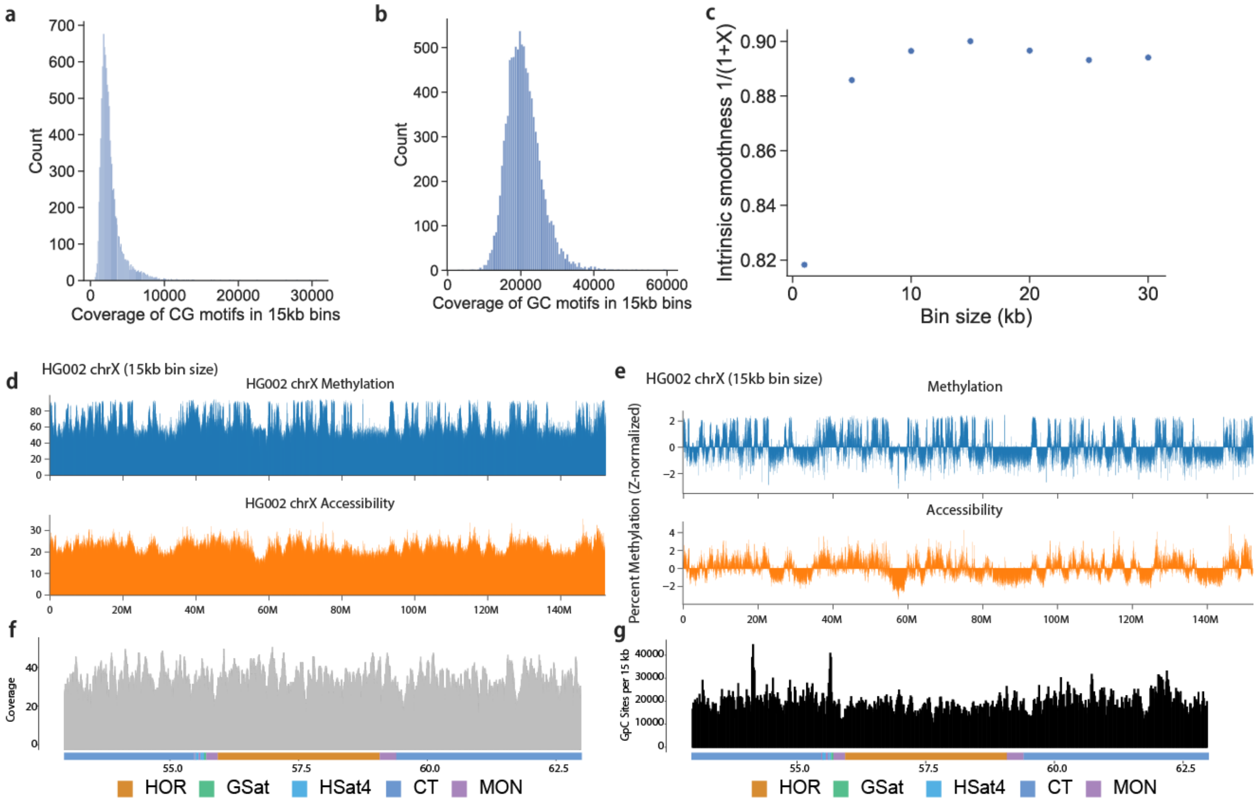


***Figure S15. NanoNOMe alignment and GpC methylation calling. a)*** *Histogram of coverage for CG sites in 15kb bins.* ***b)*** *Histogram of coverage for GC sites in 15kb bins.* ***c****) Intrinsic smoothness as a function of bin size.* ***d)*** *Top panel shows percent CpG methylation for 15kb bins across chromosome X in HG002. Bottom panel shows percent GpC methylation (accessibility) for 15kb bins across chromosome X in HG002.* ***e)*** *Top panel shows Z-normalized percent methylation (CG) for 15kb bins across the chromosome X in HG002. Bottom panel shows Z-normalized percent accessibility (GC) for 15kb bins across the chromosome X in HG002.* ***f)*** *NanoNOMe CpG call coverage in the centromeric region of the HG002 X chromosome.* ***g)*** *Number of GC sites per 15 kb bins across the centromere of HG002 chromosome X.*

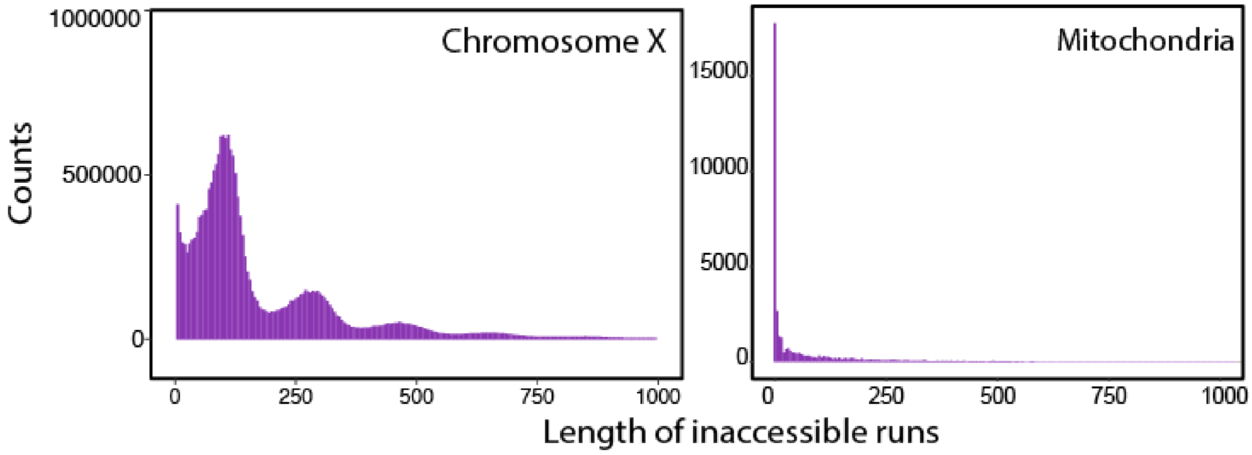


***Figure S16. NanoNOMe Inaccessible Run Lengths.*** *Histograms of the length of inaccessible runs in the NanoNOMe data. Top histogram is all of chromosome X. Peaks correspond to mono-, di-, tri- and poly-nucleosomes. Inaccessible runs occur when nucleosomes impede the ability of the GpC methyltransferase to label the DNA. Right panel shows histogram of runs on mitochondrial DNA that does not contain nucleosomes.*

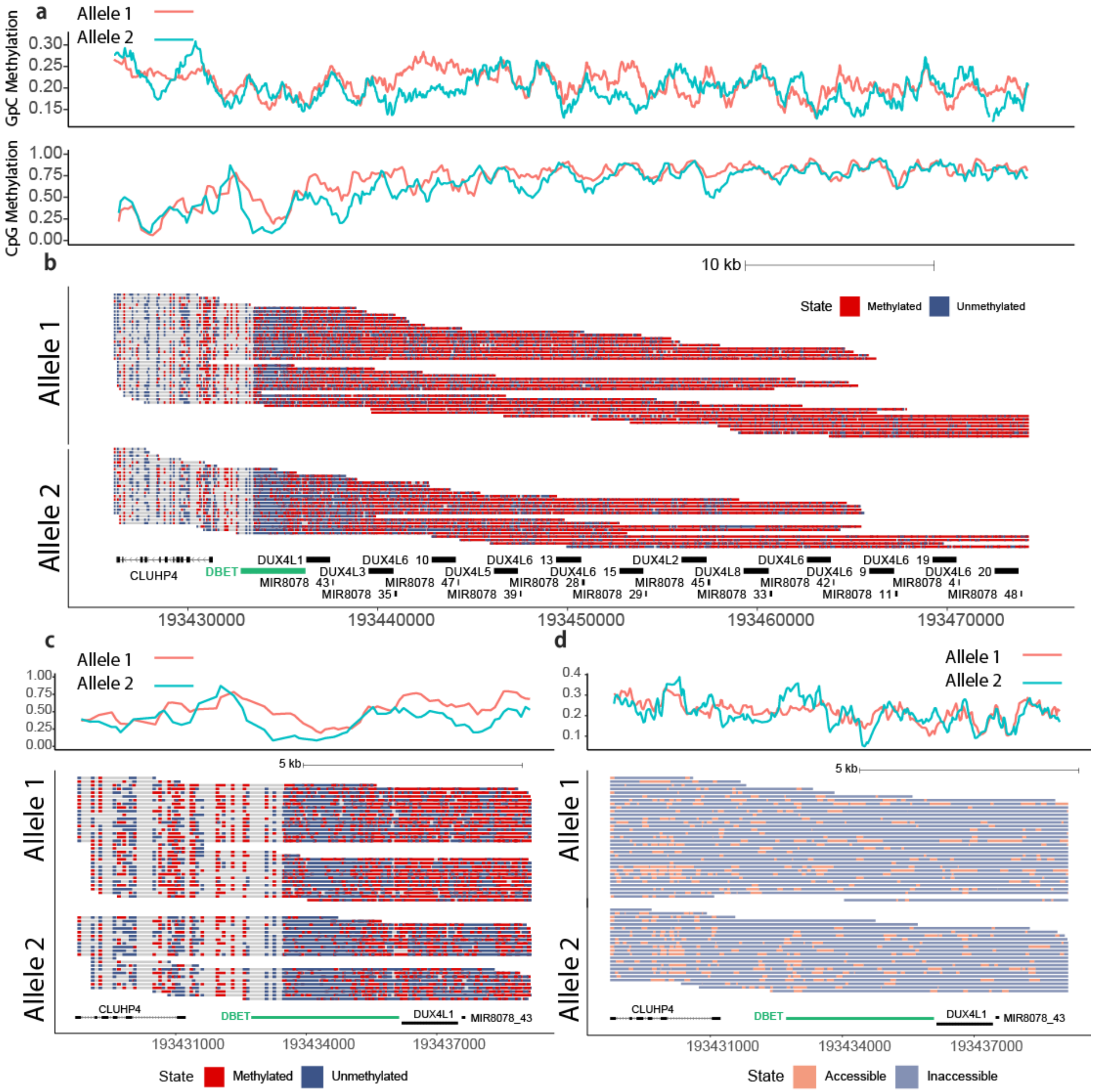


***Figure S17. Phased D4Z4 methylation frequency and accessibility in HG002. a)*** Allele specific CpG and GpC methylation at the q-subtelomere of chromosome 4. ***b)*** *Single read CpG plot of phased nanoNOMe reads at the D4Z4 locus illustrated in **a**. **c)** Single read phased methylation and **d)** accessibility at the DBET locus of the D4Z4 array.*

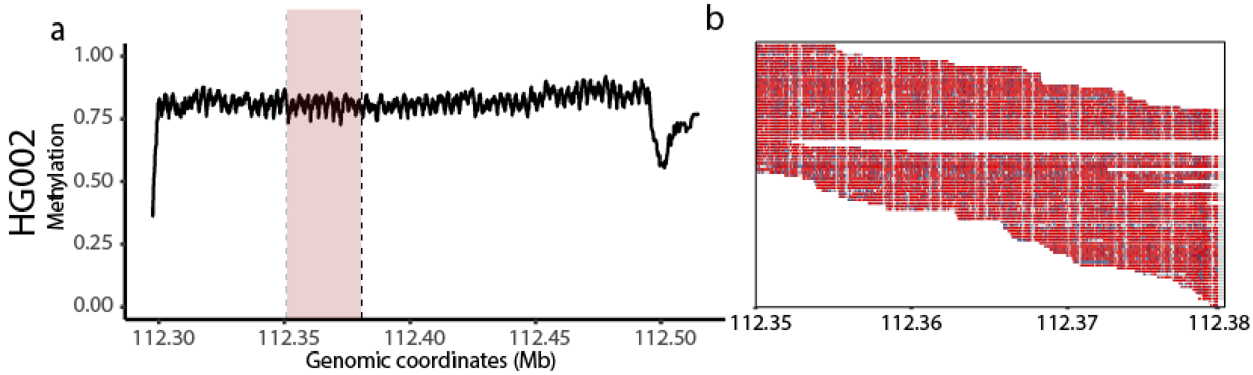


***Figure S18. DXZ4 methylation frequency in HG002. a)*** *DXZ4 methylation frequency in HG002* ***b)*** *Single read methylation plot of HG002 DXZ4 at the pink highlighted region from* ***a****.*

**Supplementary Tables:**

| Mapping Step | Target | CHM13 Mapped Reads | GRCh38 Mapped Reads | Percent Increase | Difference |
|---|---|---|---|---|---|
| mapped | Control | 1,137,935,667 | 1,131,245,094 | 0.59 | **6,690,573** |
| filtered | Control | 1,068,198,264 | 1,054,596,257 | 1.29 | **13,602,007** |
| dedup | Control | 907,959,643 | 895,616,890 | 1.38 | **12,342,753** |
| kmer | Control | 895,290,569 | 875,165,210 | 2.30 | **20,125,359** |
| mapped | CTCF | 587,838,526 | 584,410,563 | 0.59 | **3,427,963** |
| filtered | CTCF | 555,593,002 | 547,091,578 | 1.55 | **8,501,424** |
| dedup | CTCF | 440,390,633 | 433,341,439 | 1.63 | **7,049,194** |
| kmer | CTCF | 433,092,673 | 422,484,201 | 2.51 | **10,608,472** |
| mapped | H3K27ac | 1,350,669,439 | 1,346,415,081 | 0.32 | **4,254,358** |
| filtered | H3K27ac | 1,296,963,081 | 1,274,077,808 | 1.80 | **22,885,273** |
| dedup | H3K27ac | 974,695,517 | 956,778,872 | 1.87 | **17,916,645** |
| kmer | H3K27ac | 969,295,166 | 946,227,688 | 2.44 | **23,067,478** |
| mapped | H3K27me3 | 1,004,101,945 | 999,512,817 | 0.46 | **4,589,128** |
| filtered | H3K27me3 | 949,748,184 | 935,278,157 | 1.55 | **14,470,027** |
| dedup | H3K27me3 | 718,792,234 | 707,446,412 | 1.60 | **11,345,822** |
| kmer | H3K27me3 | 713,043,104 | 696,519,229 | 2.37 | **16,523,875** |
| mapped | H3K36me3 | 1,145,335,139 | 1,140,455,957 | 0.43 | **4,879,182** |
| filtered | H3K36me3 | 1,084,031,258 | 1,068,355,578 | 1.47 | **15,675,680** |
| dedup | H3K36me3 | 875,082,927 | 861,834,014 | 1.54 | **13,248,913** |
| kmer | H3K36me3 | 868,733,574 | 849,235,815 | 2.30 | **19,497,759** |
| mapped | H3K4me1 | 634,370,589 | 630,394,227 | 0.63 | **3,976,362** |
| filtered | H3K4me1 | 603,566,061 | 595,016,286 | 1.44 | **8,549,775** |
| dedup | H3K4me1 | 515,834,569 | 508,198,927 | 1.50 | **7,635,642** |
| kmer | H3K4me1 | 512,517,911 | 502,079,996 | 2.08 | **10,437,915** |
| mapped | H3K4me3 | 659,107,457 | 657,313,902 | 0.27 | **1,793,555** |
| filtered | H3K4me3 | 632,278,959 | 620,994,546 | 1.82 | **11,284,413** |
| dedup | H3K4me3 | 476,038,277 | 467,416,275 | 1.84 | **8,622,002** |
| kmer | H3K4me3 | 473,743,814 | 462,551,850 | 2.42 | **11,191,964** |
| mapped | H3K9me3 | 1,025,849,738 | 1,015,315,453 | 1.04 | **10,534,285** |
| filtered | H3K9me3 | 947,159,548 | 935,677,696 | 1.23 | **11,481,852** |
| dedup | H3K9me3 | 741,563,924 | 731,430,014 | 1.39 | **10,133,910** |
| kmer | H3K9me3 | 733,373,171 | 716,161,956 | 2.40 | **17,211,215** |

***Table S1. Statistics of dynamic k-mer assisted mapping of ENCODE data**. Table shows the total number and mean (across all cell lines) of the percentage of increase in aligned reads comparing GRCh38 and CHM13 references for each mapping step and dataset.*

| Repeat | CHM13 CpGs | GRCh38 CpGs | Difference |
|---|---|---|---|
| LINE/SINE | 10,541,816 | 10,599,859 | −58,043 |
| non−repetitive | 12,425,177 | 12,445,576 | −20,397 |
| Repeat | 4,088,225 | 3,189,158 | 899,067 |
| Satellite | 2,693,793 | 1,043,761 | 1,650,030 |
| SegDup | 2,527,456 | 1,897,529 | 629,927 |
| **Total** | **32,276,467** | **29,175,883** | **3,100,584** |

***Table S2. New CpG Regions in CHM13.*** *Number of CpG sites within different genomic regions. Difference represents the number of CpGs in GRCh38 subtracted from the number of CpGs in T2T-CHM13.*

| Repeat | Number of High Quality calls | Number of Low Quality calls | Total calls | Percent high quality |
|---|---|---|---|---|
| BSat | 3,061,606 | 1,419,809 | 4,481,415 | 68.32 |
| CT | 56,474,535 | 22,522,916 | 78,997,451 | 71.49 |
| DHOR | 646,140 | 260,432 | 906,572 | 71.27 |
| GSat | 447,699 | 184,647 | 632,346 | 70.80 |
| HOR | 29,821,344 | 11,976,265 | 41,797,609 | 71.35 |
| HSat1 | 785,693 | 350,516 | 1,136,209 | 69.15 |
| HSat2 | 23,242,685 | 7,722,744 | 30,965,429 | 75.06 |
| HSat3 | 17,244,541 | 10,358,389 | 27,602,930 | 62.47 |
| HSat4 | 72,378 | 38,109 | 110,487 | 65.51 |
| HSat5 | 41,764 | 19,356 | 61,120 | 68.33 |
| MON | 2,712,616 | 1,239,187 | 3,951,803 | 68.64 |
| **WG** | **891,974,523** | **351,636,077** | **1,243,610,600** | **71.72** |

***Table S3. Summary table of methylation high quality calls.*** *Summary statistics for CHM13 CpG calls passing LLR quality thresholds (-1.5/1.5) in each repeat type. Whole genome (WG) summary statistics are for all methylation calls from reads greater than 50 kb.*

| Tissue | Replicates | Total Reads | Accession |
|---|---|---|---|
| 1st-PB1 | . | 19054814 | SRR950976 |
| 1st-PB2 | . | 24346847 | SRR950977 |
| 1st-PB3 | . | 19926829 | SRR951019 |
| 2-cell1 | . | 18494789 | SRR950980 |
| 2-cell2 | . | 28407553 | SRR950981 |
| 2nd-PB1 | . | 27612904 | SRR950978 |
| 2nd-PB2 | . | 29538440 | SRR950979 |
| 4-cell1 | . | 49930006 | SRR950982 |
| 4-cell2 | . | 29459042 | SRR950983 |
| 8-cell1 | rep1,rep2 | 18439985 | SRR950984,SRR950985 |
| 8-cell2 | rep1,rep2 | 18411115 | SRR950986,SRR950987 |
| 8-cell3 | rep1,rep2 | 24167297 | SRR950988,SRR950989 |
| ICM1 | rep1,rep2 | 44356448 | SRR950990,SRR950991 |
| ICM2 | . | 25390088 | SRR950992 |
| ICM3 | . | 24028269 | SRR950993 |
| MII-Oocyte1 | . | 31324294 | SRR950994 |
| MII-Oocyte2 | . | 36161928 | SRR950995 |
| Morula1 | rep1,rep2 | 21366884 | SRR950996,SRR950997 |
| Morula2 | rep1,rep2 | 18252235 | SRR950998,SRR950999 |
| Morula3 | rep1,rep2 | 18454238 | SRR951000,SRR951001 |
| Postimplantation1 | rep1,rep2 | 33538771 | SRR951002,SRR951003 |
| Postimplantation2 | rep1,rep2 | 30183523 | SRR951004,SRR951005 |
| Postimplantation3 | rep1,rep2 | 27624293 | SRR951006,SRR951007 |
| Sperm1 | . | 10144406 | SRR951008 |
| Sperm2 | . | 20906531 | SRR951009 |
| Sperm3 | . | 23276912 | SRR951010 |
| Sperm4 | . | 29371875 | SRR951011 |
| TE1 | rep1,rep2 | 19269727 | SRR951012,SRR951013 |
| TE2 | rep1,rep2 | 20779347 | SRR951014,SRR951015 |
| TE3 | . | 25222928 | SRR951016 |
| Zygote1 | . | 45890346 | SRR951017 |
| Zygote2 | . | 29978135 | SRR951018 |

***Table S4. Accessions of RRBS data****. Summary of publically available RRBS data used for Figure S9 from Guo et al., 2014(2)*

| | CHM13 | | | HG002 | | | |
|---|---|---|---|---|---|---|---|
| **Repeat** | **Poor CpGs** | **Total CpGs** | **Fraction** | **Poor CpGs** | **Total CpGs** | **Fraction** | **Difference** |
| **BSat** | 1,062 | 127,898 | 0.008 | 26,966 | 127,898 | 0.211 | -0.20 |
| **CT** | 6,106 | 1,984,128 | 0.003 | 39,853 | 1,984,346 | 0.020 | -0.02 |
| **DHOR** | 4 | 20,902 | 0.000 | 207 | 20,902 | 0.010 | -0.01 |
| **Gsat** | 28 | 21,260 | 0.001 | 1,775 | 21,260 | 0.083 | -0.08 |
| **HOR** | 19,870 | 1,108,548 | 0.018 | 320,633 | 1,108,683 | 0.289 | -0.27 |
| **HSat1** | 4,905 | 30,426 | 0.161 | 11,465 | 30,426 | 0.377 | -0.22 |
| **HSat2** | 53,049 | 1,272,962 | 0.042 | 458,281 | 1,272,962 | 0.360 | -0.32 |
| **HSat3*** | 4,332 | 418,728 | 0.010 | 65,300 | 418,728 | 0.156 | -0.15 |
| **HSat4** | 1 | 2,477 | 0.000 | - | 2,479 | 0.000 | 0.00 |
| **HSat5** | - | 1,396 | 0.000 | 112 | 1,396 | 0.080 | -0.08 |
| **MON** | 38 | 85,974 | 0.000 | 1,238 | 85,965 | 0.014 | -0.01 |

*** Removal of the Chromosome 9 array**

***Table S5. CpG site coverage thresholding*****.** *Number of CpG sites with call coverage outside the coverage thresholds of 10X-100X. Poor CpGs refers to the number of CpG sites that do not meet the coverage thresholds. Total CpGs are the total number of CpGs in that repeat type. Fraction refers to the fraction of CpGs per repeat type that do not meet the coverage thresholds. The difference column is the CHM13 fraction subtracted from the HG002 fraction to show where methylation calls are of lower quality in HG002.*

| Repeat | CHM13 Methylation |
|---|---|
| BSat | 0.217 |
| CT | 0.263 |
| DHOR | 0.206 |
| Gsat | 0.158 |
| HOR | 0.56 |
| HSat1 | 0.143 |
| HSat2 | 0.176 |
| HSat3 | 0.091 |
| HSat4 | 0.115 |
| HSat5 | 0.103 |
| MON | 0.107 |
| **WG** | 0.368 |

***Table S6. CHM13 repeat median methylation values.*** *Median methylation in each repeat and for the whole genome (WG).*

| Repeat | HG002 Methylation | CHM13 Methylation |
| --- | --- | --- |
| CT | 0.818 | 0.167 |
| GSat | 0.75 | 0.308 |
| HOR | 0.654 | 0.318 |
| HSat4 | 0.522 | 0.111 |
| MON | 0.5 | 0.116 |

***Table S7. Median methylation in HG002 and CHM13 repeats on the X chromosome.*** *Median methylation in each repeat and for comparing CHM13 chromosome X and HG002 chromosome X.*

**Supplementary Data:**

**Supplemental Data 1: Summary of ENCODE data and results**. First tab (encode samples) summarizes Cell type, ChIP target and accession for datasets. Second tab (mapping stats) summarizes mapping results for different stages of the k-mer filtering approach for alignments against T2T-CHM13 or GRCh38p13. Third tab (mac2 peaks) summarizes numbers of peaks and overlap between liftover and natively called peaks.

### REFERENCES


1. J. Foox, J. Nordlund, C. Lalancette, T. Gong, M. Lacey, S. Lent, B. W. Langhorst, V. K. C. Ponnaluri, L. Williams, K. Padmamabhan, R. Cavalcante, A. Lundmark, D. Butler, J. Gurvitch, J. M. Greally, M. Suzuki, M. Menor, M. Nasu, A. Alonso, C. Sheridan, A. Scherer, S. Bruinsma, G. Golda, A. Muszynska, P. P. Labaj, M. A. Campbe, F. Wos, A. Raine, U. Liljedahl, T. Axelsson, C. Wang, Z. Chen, Z. Yang, J. Li, X. Yang, H. Wang, A. Melnick, S. Guo, A. Blume, V. Franke, I. Ibanez de Caceres, C. Rodriguez-Antolin, R. Rosas, J. Wade Davis, J. Ishii, D. B. Megherbi, W. Xiao, W. Liao, J. Xu, H. Hong, B. Ning, W. Tong, A. Akalin, Y. Wang, C. E. Mason, The SEQC2 Epigenomics Quality Control (EpiQC) Study: comprehensive characterization of epigenetic methods, reproducibility, and quantification. *bioRxiv* (2020), , doi:10.1101/2020.12.14.421529.

2. H. Guo, P. Zhu, L. Yan, R. Li, B. Hu, Y. Lian, J. Yan, X. Ren, S. Lin, J. Li, X. Jin, X. Shi, P. Liu, X. Wang, W. Wang, Y. Wei, X. Li, F. Guo, X. Wu, X. Fan, J. Yong, L. Wen, S. X. Xie, F. Tang, J. Qiao, The DNA methylation landscape of human early embryos. *Nature*. **511**, 606–610 (2014).
